# Age-linked lung pathology is reduced by immunotherapeutic targeting of isoDGR protein damage

**DOI:** 10.1101/2023.08.11.552994

**Authors:** Pazhanichamy Kalailingam, Khalilatul-Hanisah Mohd-Kahliab, SoFong Cam Ngan, Ranjith Iyappan, Benjamin Sian Teck Lee, Bhargy Sharma, Radek Machan, Sint Thida Bo, Emma S Chambers, Val A Fajardo, Rebecca E. K. Macpherson, Jian Liu, Panagiota Klentrou, Evangelia Litsa Tsiani, Kah Leong Lim, I Hsin Su, Yong-Gui Gao, A Mark Richar, Raj N. Kalaria, Christopher P. Chen, Neil E. McCarthy, Siu Kwan Sze

## Abstract

Advancing age is the primary risk factor for pulmonary diseases. Our investigation revealed an 8-fold increase in aging induced isoDGR-damaged proteins in lung tissue from human pulmonary fibrosis patients compared to healthy tissues, accompanied by elevated frequencies of CD68+/CD11b+ macrophages, indicating lung tissue is susceptible to time-dependent accumulation of isoDGR-proteins. To elucidate the mechanisms through which isoDGR-proteins may exacerbate aging lung disorders for potential therapeutic targeting, we assessed the functional role of this isoDGR-motif in naturally-aged mice and mice lacking the corresponding isoDGR repair enzyme (Pcmt1-/-). IsoDGR-protein accumulation in mouse lung tissue and blood vessels correlated with chronic low-grade inflammation, pulmonary edema, and hypoxemia. IsoDGR accretion induced mitochondrial and ribosomal dysfunctions, cellular senescence, and apoptosis, contributing to progressive lung damage over time. Treatment with anti-isoDGR antibodies suppressed TLR pathway activity, mitigated cytokine-driven inflammation, restored mtDNA expression, and significantly reduced lung pathology in-vivo. Similarly, exposure of lung endothelial cells to isoDGR-modified fibronectin impaired oxygen consumption, increased reactive oxygen species levels, and disrupted acidification, but these effects were efficiently reversed by target-specific antibody therapy. Collectively, our findings underscore the significant contribution of isoDGR-damaged proteins to age-linked lung pathology. IsoDGR-specific therapy emerges as a promising treatment approach for pulmonary disorders in older patients.

## Introduction

Advancing age is the primary risk factor for significant lung pathology and associated mortality^1^. Numerous studies have highlighted that pulmonary function, crucial for supplying oxygen to tissues, strongly predicts human morbidity and mortality^2–7^. Human lungs exhibit the largest surface area in the body, comprising hundreds of millions of alveolar sacs and an extensive capillary blood vessel network that facilitate rapid oxygen exchange^1^. The lung is supported by a scaffold formed of extracellular matrix (ECM) components and resident interstitial cells that regulate pulmonary functions and local immune responses, including epithelial cells, alveolar macrophages (AMs), and peribronchial interstitial macrophages (IMs)^1^. Pulmonary ECM proteins exhibit slow turnover and are therefore susceptible to accumulation of molecular damage that contributes to lung aging^8^ including structural and functional changes such as loss of elasticity, increased tissue inflammation, and reduced muscle strength. Accordingly, dysregulation of pulmonary ECM turnover has been shown to induce chronic inflammation of the airways and been implicated in the development of respiratory diseases^9^ including emphysema, chronic obstructive pulmonary disease (COPD), asthma, fibrosis, and acute respiratory distress syndrome^1,10,11^.

At a fundamental level, aging results from damage to key biomolecules via accumulation of degenerative protein modifications (DPMs) *in vivo*, leading to the gradual deterioration of tissue function and increased vulnerability to disease and death^1,12–17^. We previously reported that age-linked damage to the amino acid sequence NGR (Asn-Gly-Arg) results in a ‘gain-of-function’ conformational switching to isoDGR motifs (isoAsp-Gly-Arg) that can mediate integrin binding^12,18–20^. We observed that extracellular isoDGR accumulates in body tissues from elderly human patients and in animal models of aging, while functional experiments confirmed that this motif can activate macrophages to promote chronic inflammation. Accordingly, immunotherapy directed against isoDGR was sufficient to trigger immune clearance of isoDGR-damaged proteins, reduce inflammatory cytokine levels, and double the average lifespan of mice that lack the corresponding repair enzyme Pcmt1^21–23^. However, the specific tissues and organ systems most impacted in this context have yet not been identified, and the physiological relevance to elderly human patients has remained unclear.

IsoDGR modification has previously been observed in structural proteins fibronectin, laminin, tenascin C, and several other ECM constituents of human artery, leading to increased leukocyte infiltration of coronary vessels^19,20^. These ECM proteins are also integral components of human lungs, which form a complex anatomy of fibrous proteins (collagen, elastin), glycoproteins (fibronectin, laminin), glycosaminoglycans (heparin, hyaluronic acid) and proteoglycans (perlecan, versican)^8^. These long-lived lung proteins are particularly susceptible to isoDGR accumulation, and may trigger macrophage infiltration and express pro-inflammatory cytokines^20^. Therefore, the isoDGR-damaged ECM may induce pulmonary ‘inflamm-aging’ and promote age-linked lung disease. In the current study, we hypothesized that age-associated deposition of isoDGR in lung ECM activates recruited/resident monocyte-macrophages, thereby promoting low-grade inflammation and eventual pulmonary dysfunction. To test this hypothesis, we examined isoDGR accumulation via immunostaining of human lung tissues from healthy individuals and patients with lung fibrosis, which revealed an 8-fold increased level of isoDGR-protein levels in fibrotic tissues, accompanied by marked infiltration of CD68+/CD11b+ macrophages. We next investigated the mechanistic link between isoDGR accumulation and age-associated pulmonary disease in both naturally-aged mice and animals that lack the isoDGR repair enzyme Pcmt1. We observed marked accumulation of isoDGR-modified proteins in murine lungs that was accompanied by pulmonary inflammation, enlarged airspaces, tissue oedema, congested blood vessels, and pronounced hypoxemia. Transcriptomic and functional analyses further showed that isoDGR dysregulates ribosome activity, electron transport chain (ETC), and core mitochondrial functions, while also promoting generation of reactive oxygen species (ROS) and driving senescence / apoptosis of lung cells. Conversely, treatment with isoDGR-specific antibody successfully reduced lung pathology by decreasing TLR pathway activity and tissue inflammation, in addition to reducing redox stress and mitochondrial dysfunction *in vivo*.

## Results

### Advancing age and lung fibrosis are associated with isoDGR and macrophage accumulation

We first aimed to assess isoDGR-modified protein levels in aging lung and fibrotic pulmonary tissue to investigate potential associations with CD68+/CD11b+ macrophage infiltration. To do this, we acquired two human lung tissue microarrays (TMA-LC2086a and TMA-LC561; from Tissue Array Technology USA) comprising 192 sections of healthy human lung tissue and 56 sections of pulmonary interstitial fibrosis tissue. Analysis immunohistochemistry confirmed that isoDGR-modified protein levels in normal human lung tissues are increased with advanced age, regardless of genetic background (isoDGR staining was positively correlated with age by linear regression; slope 3.33). We also detected CD68+ and CD11b+ cells in most of the aging human lung tissue samples, and frequencies were positively correlated with isoDGR levels. These findings clearly suggest that isoDGR-modified protein damage increases with aging and is associated with immune cell infiltration, which may contribute to a range of age-linked lung disorders (Fig.1). Indeed, in a separate analysis of tissues from pulmonary fibrosis cases and lung cancer patients (Fig. S1), we observed that age-related accumulation of isoDGR-proteins was increased 8-fold increase in disease settings (slope 26.20 by linear regression). Again, isoDGR levels were positively correlated with localization of CD68+ and CD11b+ immune cells, indicating that isoDGR-motif may contribute to macrophage activation and pro-fibrotic lung inflammation.

**Figure 1:**
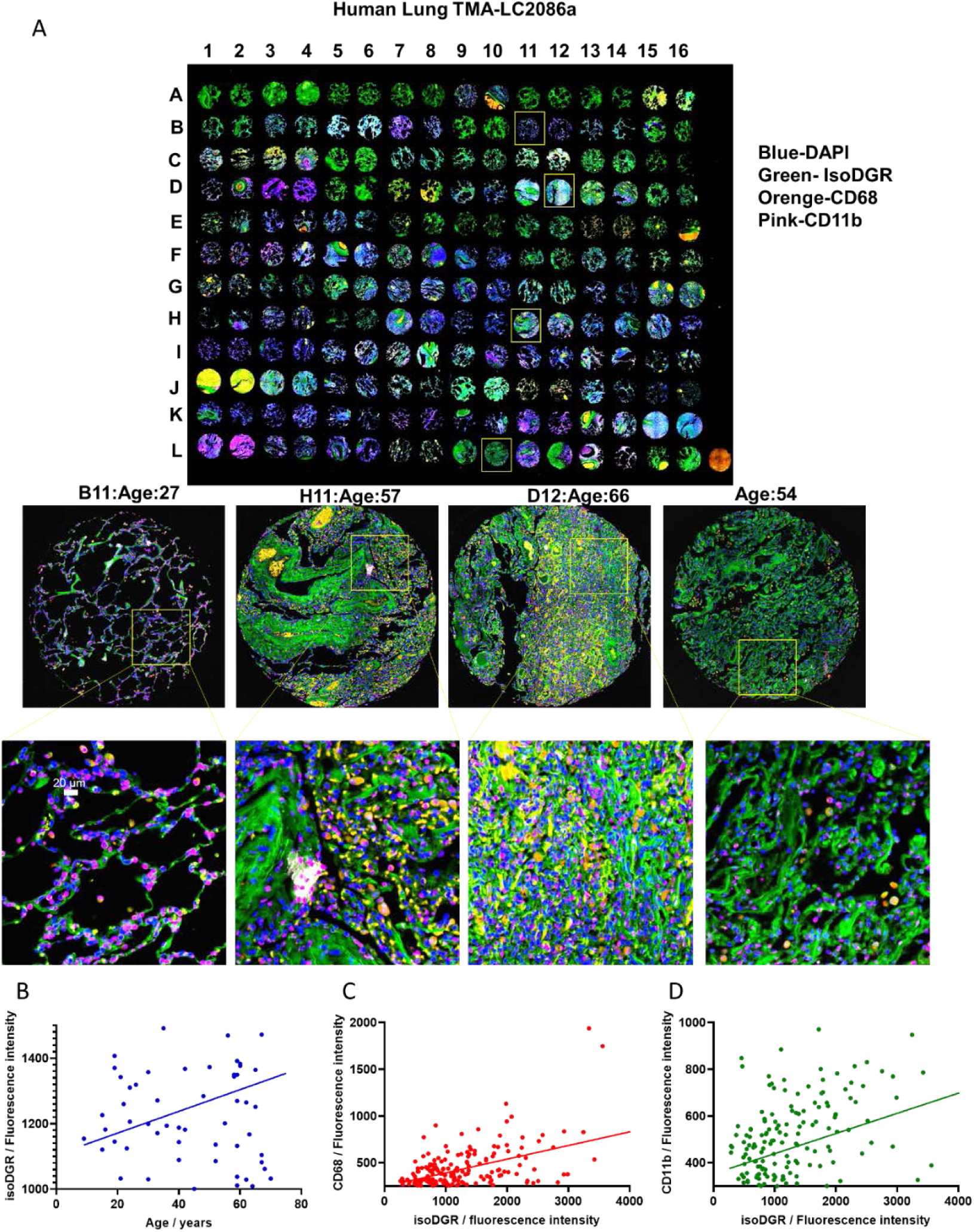
Age-induced accumulation of isoDGR-modified proteins correlates with CD68+ and CD11b+ cells in human lung tissue. (A) Representative immunostaining images showing isoDGR-protein distribution and correlation with CD68+ and CD11b+ immune cells in 192 sections of normal human lung tissues of varying age. Representative zoomed images are shown; B11: Age 27, H11: Age57, D12: Age 66, L10: Age 54. (B) IsoDGR level was positively correlated with age (linear regression slope 3.33). (C) CD68 level was positively correlated with isoDGR level. (D) CD11b level was positively correlated with isoDGR level.

### IsoDGR protein damage promotes age-linked lung pathology

We next sought to confirm that isoDGR-modified proteins can promote age-linked lung pathology and assess whether this motif represents a promising target for novel immunotherapies. For this, we investigated the functional role of isoDGR in both naturally-aged mice and animals that lack the corresponding repair enzyme Pcmt1. Deletion of Pcmt1 results in extensive accumulation of isoDGR protein damage in mouse body tissues^24^, leading to premature aging and a reduced lifespan of just 4-12 weeks^25–27^. In previous work, we showed that isoDGR interacts with macrophage integrins to induce pathological release of TNFα and MCP1^20^, whereas weekly treatment with 1mg/kg isoDGR-specific mAb reduced vascular inflammation and doubled the lifespan of Pcmt1^-/-^ mice^12^. To investigate the pathological effects of isoDGR accretion across different organs, in this study we first generated global Pcmt1^-/-^ mice by crossing Pcmt1^+/-^ males with Pcmt1^+/-^ females (genotype was confirmed by PCR of tail genomic DNA; 25% of pups born were Pcmt1^-/-^ consistent with the expected Mendelian ratio). Pcmt1 deletion in mouse lung tissue was confirmed by western blot of protein extract from knockout and wild-type animals (Fig 2A). Further western blot analysis indicated that isoDGR-modified lung proteins accumulated to high levels in Pcmt1^-/-^ mice compared with WT control mice, and isoDGR-specific mAb treatment reduced damaged protein levels in body tissues (Fig 2B & C). Examination on autopsy revealed severe pulmonary oedema in mice that died at 4-6 weeks (10%), moderate pathology in mice that died at 6-10 weeks (60%), and less severe histology in mice that survived beyond 10 weeks (30%). Moreover, accumulation of isoDGR-damaged proteins was also observed in naturally-aged WT mice (17 months old) and motif levels were reduced by target-specific mAb immunotherapy but not isotype-matched antibody treatment. (Fig. S2).

**Figure 2:**
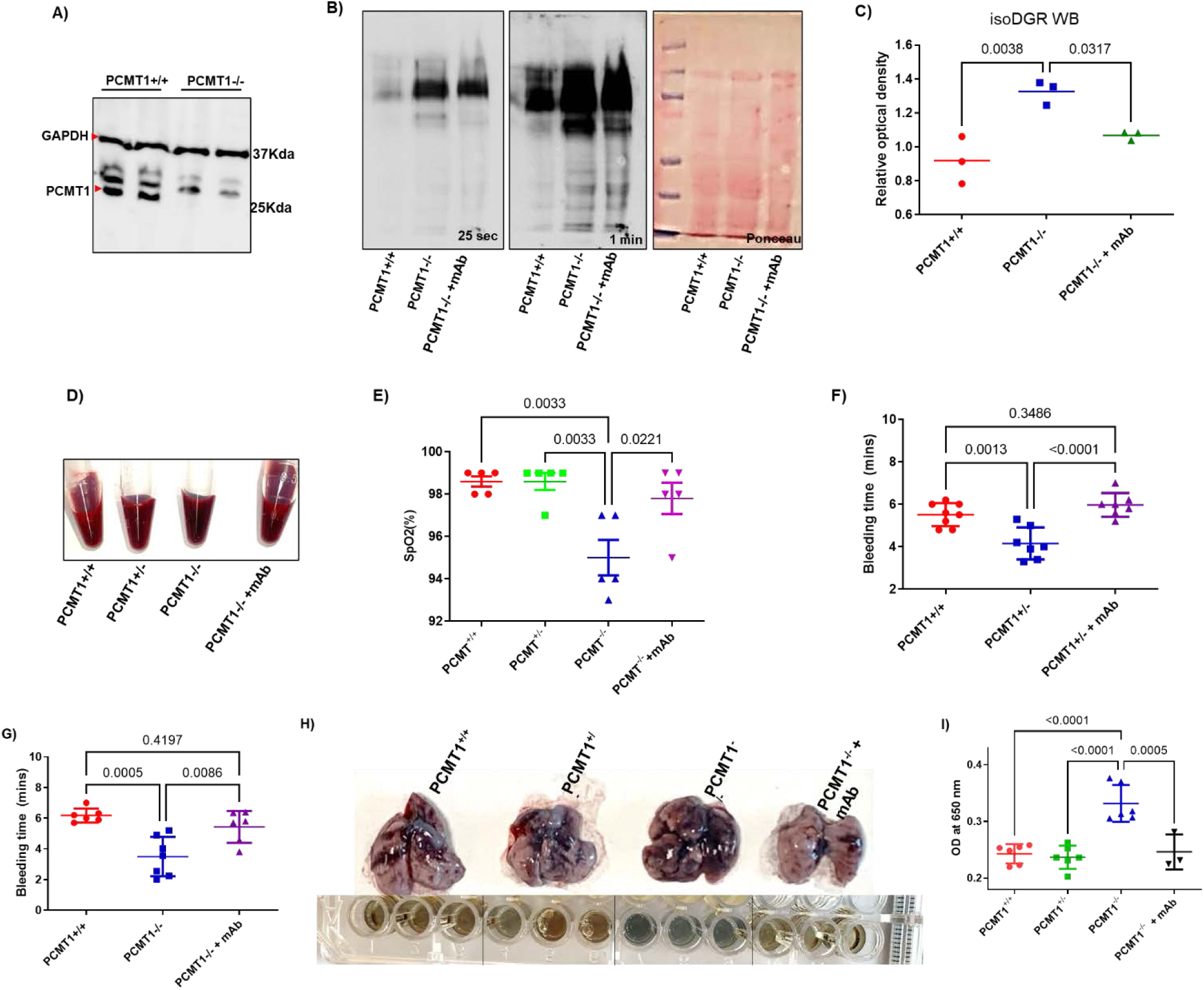
IsoDGR-damaged protein accumulation in lungs impacts oxygen exchange. **(A)** Lung protein lysates from Pcmt1^+/+^ and Pcmt1^-/-^ mice were subjected to western blot using antibodies against Pcmt1 enzyme. GAPDH was used as a loading control (n=3). **(B)** Lung protein lysates from Pcmt1^+/+^, Pcmt1^-/-^ and mAb-treated Pcmt1^-/-^ mice were subjected to western blot using isoDGR-specific mAb. Protein loading was visualized by Ponceau S.(n=3) **(C)** Graph showing quantification of isoDGR-damaged protein levels in the lungs of Pcmt1^+/+^, Pcmt1^-/-^, and mAb-treated Pcmt1^-/-^ mice. **(D)** Representative images of blood samples from each phenotype collected at 6 weeks. Blood from Pcmt1^-/-^ mice appeared dark red in colour, unlike that of mAb-treated Pcmt1^-/-^ mice. **(E)** Dot plot shows SpO2 level in WT, Pcmt1^+/-^, Pcmt1^-/-^, and mAb-treated Pcmt1^-/-^ mice at 6 weeks (n=5). SpO2 level was reduced in Pcmt1^-/-^ mice and markedly improved by mAb treatment. **(F)** Dot plot of tail bleeding time in Pcmt1^+/+^, Pcmt1^+/-^, and mAb-treated Pcmt1^+/-^ mice (n=7-8) at 1.5 years. **(G)** Dot plot of tail bleeding time in Pcmt1^+/+^, Pcmt1^-/-^, and mAb-treated Pcmt1^-/-^ mice at 6 weeks (n=6). **(H)** Representative lung appearance upon assessment of pulmonary vascular permeability by Evans blue dye extravasation assay. **(I)** Dot graph showing quantitative analysis of Evans blue-labelled albumin extravasation from the lungs of Pcmt1^+/+^ (n=6), Pcmt1^+/-^ (n=6), Pcmt1^-/-^ (n=6) and mAb-treated Pcmt1^-/-^ mice (n=3) assessed at 6 weeks. Results are shown as mean ± SEM.

### IsoDGR-induced tissue damage impairs pulmonary oxygen exchange

Having observed pulmonary oedema in Pcmt1^-/-^ mice, we next performed haematology examinations to better determine the mechanisms underlying isoDGR-induced lung pathology. Blood from Pcmt1^-/-^ and older Pcmt1^+/-^ animals was immediately noted as being darker in shade than that of Pcmt1^+/+^ mice, indicating marked accumulation of carboxyhemoglobin (Fig.2D). We therefore proceeded to check blood oxygen (SpO2) levels in Pcmt1^+/-^, Pcmt1^-/-^, mAb-treated Pcmt1^-/-^ mice, and Pcmt1^+/+^ controls. These data indicated a significant reduction of SpO2 in Pcmt1^-/-^ mice (94%-97%) compared with Pcmt1^+/+^ control mice (97%-99%), suggesting marked hypoxemia in animals that lack the isoDGR repair enzyme (Fig.2E). Given that this pathology can arise from either impaired lung function or haematological defects, we next performed a complete blood count to evaluate the likely cause of hypoxemia. RBC counts and haemoglobin levels were not significantly different between Pcmt1^-/-^ and Pcmt1^+/+^ mice, suggesting that these parameters did not play an important role in the observed hypoxemia (Fig. S3). Having previously identified that isoDGR-modified fibrinogen may promote coagulation in blood from cardiovascular disease (CVD) patients, we next performed tail bleeding assays in Pcmt1^+/-^ and Pcmt1^+/+^ mice at 1.5 years of age (by which time isoDGR-modified protein levels are significantly increased in Pcmt1^+/-^ compared to Pcmt1^+/+^ animals). Intriguingly, isoDGR accumulation was found to be associated with pro-thrombotic phenotypes, as demonstrated by a significantly reduced bleeding time in Pcmt1^+/-^ mice compared with age-matched Pcmt1^+/+^ mice (Fig.2F). Next, we performed a bleeding assay in Pcmt1^+/+^, Pcmt1^-/-^, and mAb-treated Pcmt1^-/-^ mice. We observed a significantly shorter bleeding time in Pcmt1^-/-^ mice compared to Pcmt1^+/+^ animals. Treatment with isoDGR motif-specific mAb restored bleeding time to levels comparable with Pcmt1^+/+^ mice. These results suggest that excess isoDGR-protein in body tissues is associated with pro-thrombotic phenotypes in Pcmt1^-/-^ mice. We therefore proceeded to test extravasation of Evans blue dye from the pulmonary blood vessels of Pcmt1^+/+^, Pcmt1^+/-^, Pcmt1^-/-^, and mAb-treated Pcmt1^-/-^ mice. We observed a marked increase in Evans blue extravasation from the lungs of Pcmt1^-/-^ mice, indicating significant vascular leakage consistent with pulmonary oedema, impaired oxygen exchange, and resultant hypoxemia. Crucially, isoDGR-specific mAb treatment restored blood samples to a bright red colour typical of Pcmt1^+/+^ oxygenation levels (Fig.2D), and normalised SpO2 values (Fig.2E) while also reducing coagulopathy (Fig.2F-G) and minimising vascular leakage (Fig.2H-I). Together, these results indicate that isoDGR-induced lung pathology and coagulopathy are the primary causes of hypoxemia in Pcmt1^-/-^ mice, and that isoDGR-specific immunotherapy can ameliorate these pathological features *in vivo*.

### IsoDGR-enriched lung tissues display histological features of age-linked pathology

We next performed H&E staining to better visualise lung histology in Pcmt1^+/+^, Pcmt1^+/-^, Pcmt1^-/-^, and mAb-treated Pcmt1^-/-^ mice. These analyses revealed enlargement of the perivascular space, blood vessel congestion, airspace distension, and increased perivascular leukocyte infiltration in Pcmt1^-/-^ mouse lungs (Fig.3). When lung phenotypes were assessed by MRI, we observed increased opacity and consolidation in Pcmt1^-/-^ and 2-year-old Pcmt^+/-^ mice relative to Pcmt1^+/+^ controls, consistent with significant pulmonary inflammation and oedema (Fig. S3). Pathological assessment also confirmed severe alveolar oedema that likely contributed to the premature death of Pcmt1^-/-^ mice (10% very severe oedema, 50-60% moderate, 30% less severe). Several previous reports have shown that reduced SpO2 levels in mice of advanced age is due to enlargement of alveoli, leading to impaired gas exchange with the circulation. Strikingly, weekly treatment with isoDGR-specific mAb was able to limit airspace enlargement and improved lung histology in 5-6 week-old Pcmt1^-/-^ mice (Fig.3).

**Figure 3:**
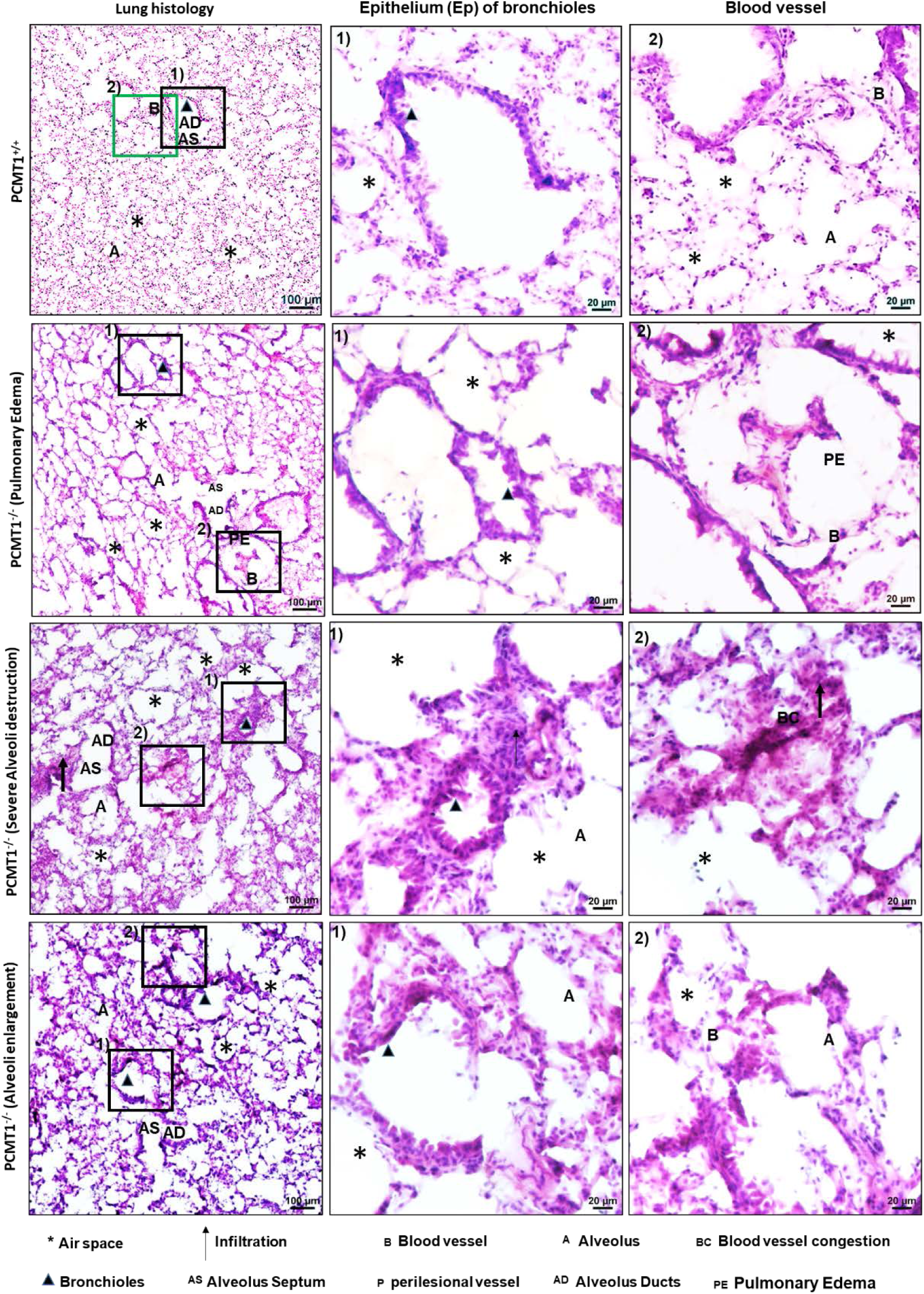
IsoDGR mAb treatment improves emphysema phenotype in lungs from Pcmt^-/-^ mice. Representative H&E-stained lung sections from 5-6 week-old Pcmt1^+/+^, Pcmt1^+/-^ and Pcmt1^-/-^ mice, displaying airspace enlargement, severe pulmonary oedema, and alveolar destruction, except in Pcmt1^-/-^ mice treated with anti-isoDGR mAb therapy. Magnified images show representative lung oedema, blood vessel congestion, epithelial (Ep) swelling in bronchioles, and leukocyte infiltration in untreated Pcmt1^-/-^ animals (n=9).

### Positive correlation of CD68+ lung leukocytes with isoDGR-modified extracellular matrix

Immunostaining of lung tissues revealed marked accumulation of isoDGR-damaged proteins in the lung parenchyma and perivascular walls of Pcmt1^-/-^ mice compared with control mice (Fig.4). Intriguingly, isoDGR distribution appeared to coincide with CD68^+^ leukocyte infiltration (Fig.4 A-C), suggesting that this motif may promote monocyte-macrophage recruitment to damaged ECM components. Indeed, the extent of isoDGR accumulation in lung tissue was found to be positively correlated with CD68^+^ monocyte-macrophage localisation (Fig.4D). Subsequent immunostaining revealed that isoDGR distribution also correlated with F4/80^+^ cells (Fig. S5) and fibronectin in the lungs of Pcmt1^-/-^ mice (Fig. S6), further suggesting that isoDGR-modified ECM components can recruit monocyte-macrophages to induce pulmonary inflammation. Importantly, mAb treatment reduced isoDGR-modified protein levels in the lungs of Pcmt1^-/-^ mice, suggesting that immune-mediated clearance of this damage motif can reduce age-linked tissue inflammation.

**Figure 4:**
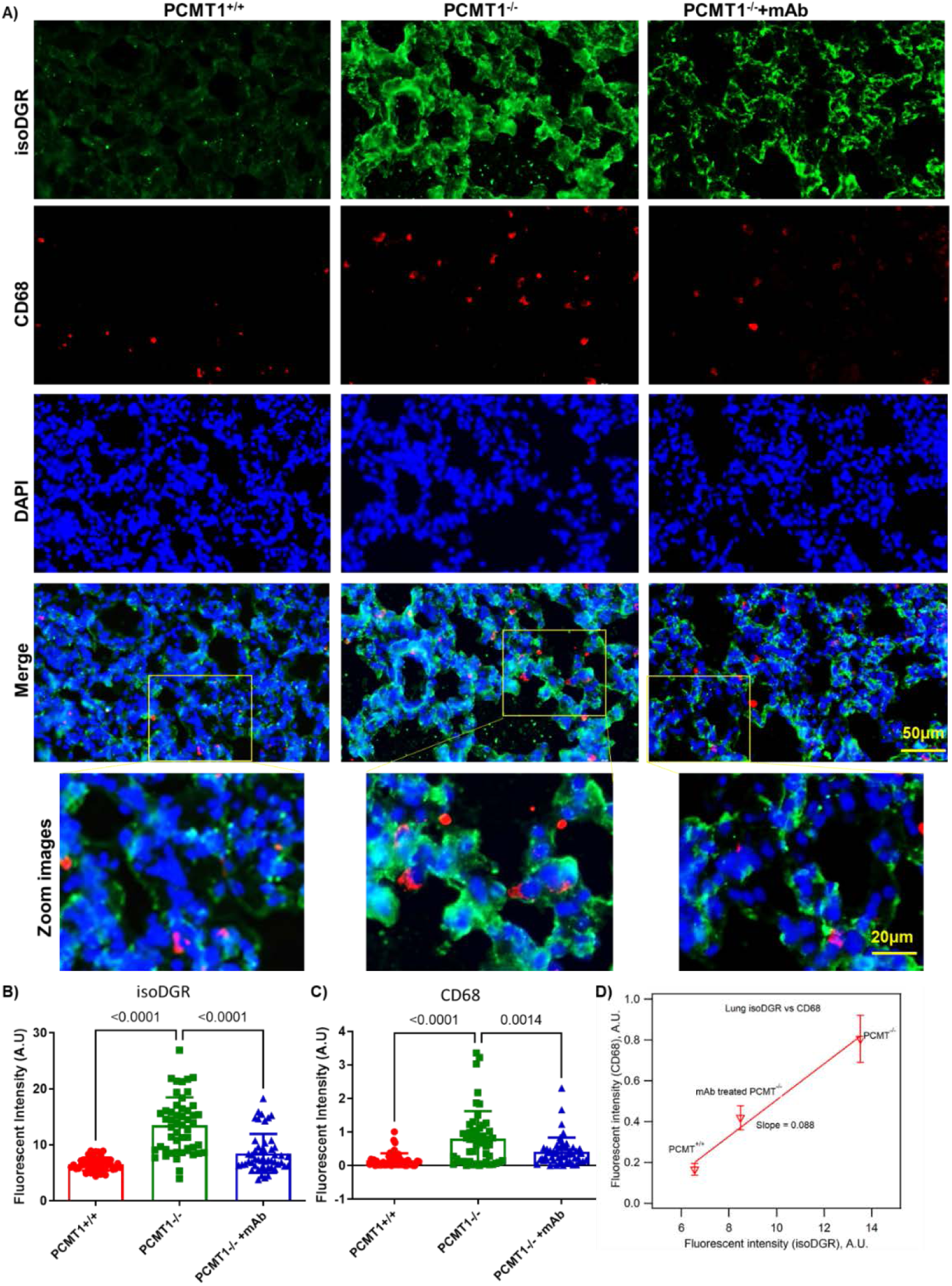
IsoDGR accumulation and positive correlation with CD68+ cell infiltration of Pcmt1^-/-^ mouse lung. **A)** Representative immunostaining of isoDGR-damaged protein distribution and correlation with CD68^+^ macrophages in cryosectioned lung tissue from Pcmt1^+/+^, Pcmt1^-/-^, and mAb-treated Pcmt1^-/-^ mice at age 5-6 weeks (n=5). IsoDGR **(B)** and CD68 **(C)** fluorescence were quantified in Image J using 50 randomized regions in 5 lung images from 5 individual mice for each genotype (graphs show mean values for the same region from 5 images). **(D)** Fluorescence intensity of CD68 was proportional to isoDGR intensity, indicating that motif distribution was associated with macrophage infiltration. Results are shown as mean values ± SEM.

### Antibody therapy reverses age-linked increases in isoDGR-mediated lung inflammation

H&E imaging of lung pathology revealed a correlation of disease severity with increasing age of Pcmt1^+/+^ and Pcmt1^+/-^ mice, although heterozygous animals displayed more severe features at the same age (assessed at 4, 15, and 24 months, Fig. S7). To confirm this apparent link between age-induced isoDGR accumulation and lung pathology, we next determined motif expression levels and infiltration of CD68+ macrophages in lung tissues from Pcmt1^+/+^ and Pcmt1^+/-^ mice aged 4, 15, or 24 months (Fig.5A-C). Analysis by lung tissue immunostaining confirmed that isoDGR- damaged protein levels and CD68+ cell frequency increased with age in both Pcmt1^+/+^ and Pcmt1^+/-^ mice (Fig.5D, E), and that Pcmt1^+/-^ mice displayed more extensive motif accumulation. Notably, CD68 fluorescence was found to increase in proportion with isoDGR intensity, and both features accumulated faster in lungs from older mice irrespective of genotype (Fig.5F). These results strongly suggest that aging leads to accumulation of isoDGR-damaged lung proteins that likely recruit CD68+ macrophages to perivascular and parenchymal tissues, thus creating a local inflammatory response that drives pulmonary pathology. We therefore tested whether age-damaged protein levels can also be reduced in the lungs of naturally-aged mice treated with target-specific immunotherapy. Indeed, we observed that injection of isoDGR-specific mAb into 17 month-old mice resulted in marked immune clearance of damaged lung proteins, whereas administration of an isotype-matched IgG control antibody did not (Fig. S8).

**Figure 5:**
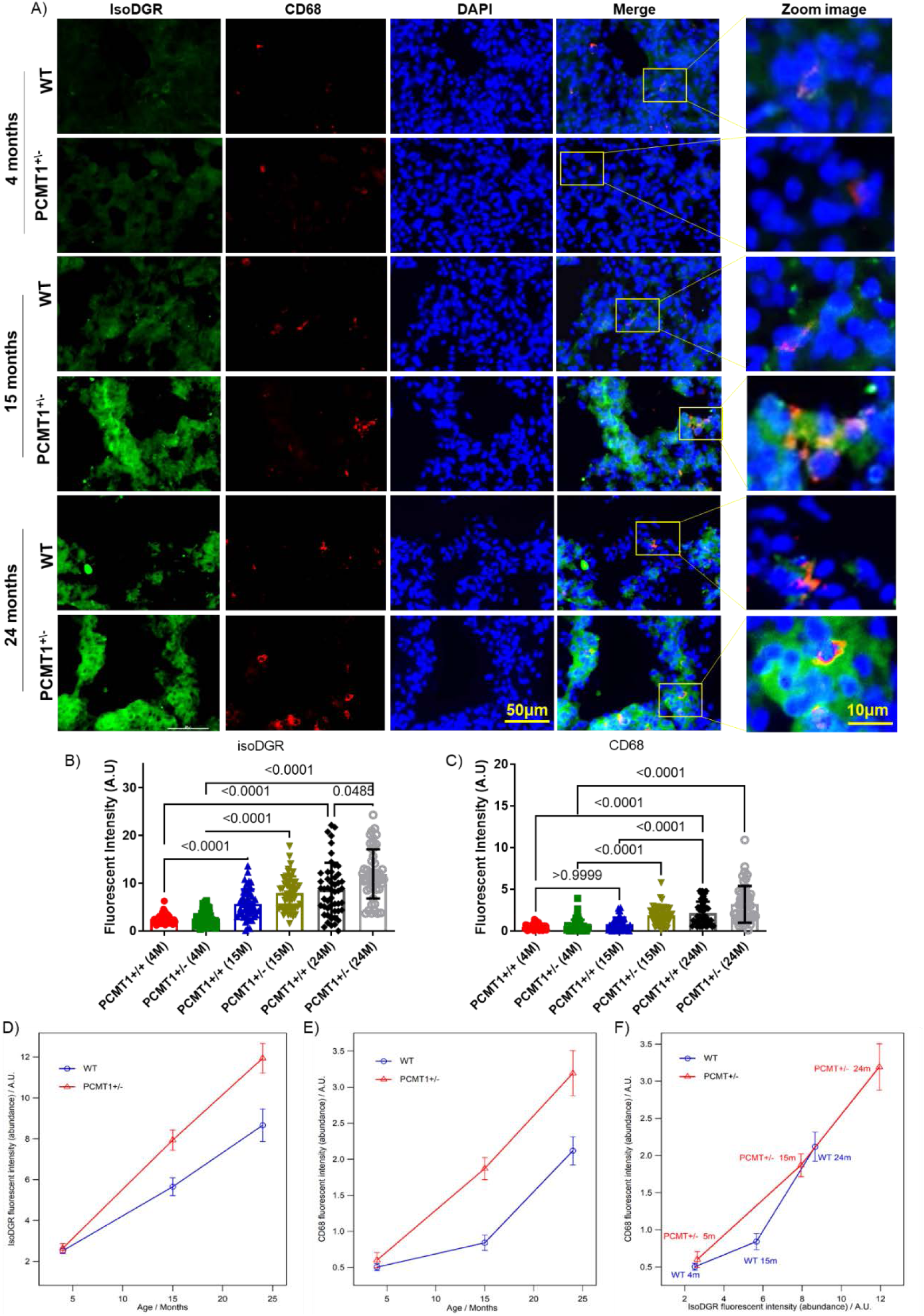
Age-induced accumulation of isoDGR-modified proteins correlates with CD68+ lung leukocytes in Pcmt1^+/-^ mice. **(A)** Representative immunostaining images showing isoDGR-protein distribution and correlation with CD68^+^ macrophages in cryosectioned lung tissue from Pcmt1^+/+^, Pcmt1^+/-^, Pcmt1^-/^,^-^ and mAb-treated Pcmt1^-/-^ mice at 4, 15, or 24 months (n=3). isoDGR **(B)** and CD68 **(C)** fluorescence were quantified in Image J using 50 randomized regions from 3 images of 3 independent lung sections for each genotype (graphs show average values for the same region from 3 images). **(D)** Plot showing age-linked accumulation of isoDGR-motif in both Pcmt1^+/+^ and Pcmt1^+/-^ animals. **(E)** Graph showing age-linked accumulation of CD68^+^ cells in both Pcmt1^+/+^ and Pcmt1^+/-^ animals. Results indicate that CD68+ macrophage infiltration is accelerated in old animals irrespective of genotype **(F)**, with cell frequency also being proportional to isoDGR levels. Statistical significance was determined by Kruskal- Wallis test. Results shown are mean values ± SEM.

### Aging-damaged isoDGR-proteins induce both local and systemic inflammatory responses

We previously reported that isoDGR binding to macrophage surface integrins induces pro-inflammatory cytokine release in artery walls, whereas others have observed that CD68+ alveolar macrophages contribute to severe pulmonary inflammation in emphysema / COPD^28^. Having already identified that resident macrophages co-localise with isoDGR-damaged lung proteins (Fig. 4, 5 and S4), we next investigated whether this association could result in increased tissue inflammation. To do this, we performed reverse transcription-quantitative polymerase chain reaction (RT-qPCR) using total RNA extracted from the lungs of Pcmt1^+/+^, Pcmt1^+/-^, Pcmt1^-/-^, and mAb-treated Pcmt1^-/-^ mice. Using this analysis, lungs from Pcmt1^-/-^ mice were found to display significantly higher expression of inflammatory mediators including MCP1, IL-1α IL-12p40, IL-8, CCL4 and IL-3. Strikingly, mAb treatment of Pcmt1^-/-^ mice reduced lung tissue levels of these same inflammatory cytokines while simultaneously increasing expression of pro-regulatory IL-10 (Fig.6). These results suggest that motif-specific immunotherapy can promote antibody-mediated phagocytic clearance of isoDGR-damaged lung proteins to reduce pulmonary inflammation^12^. Similarly, when we quantified inflammatory cytokines in 2-year-old mouse lungs, we detected increased tissue levels of MCP1, TNFα and IL-1α in Pcmt1^+/-^ mice compared with age-matched Pcmt1^+/+^ controls (Fig. S9A), consistent with age-linked patterns of isoDGR accumulation in these genotypes.

**Figure 6:**
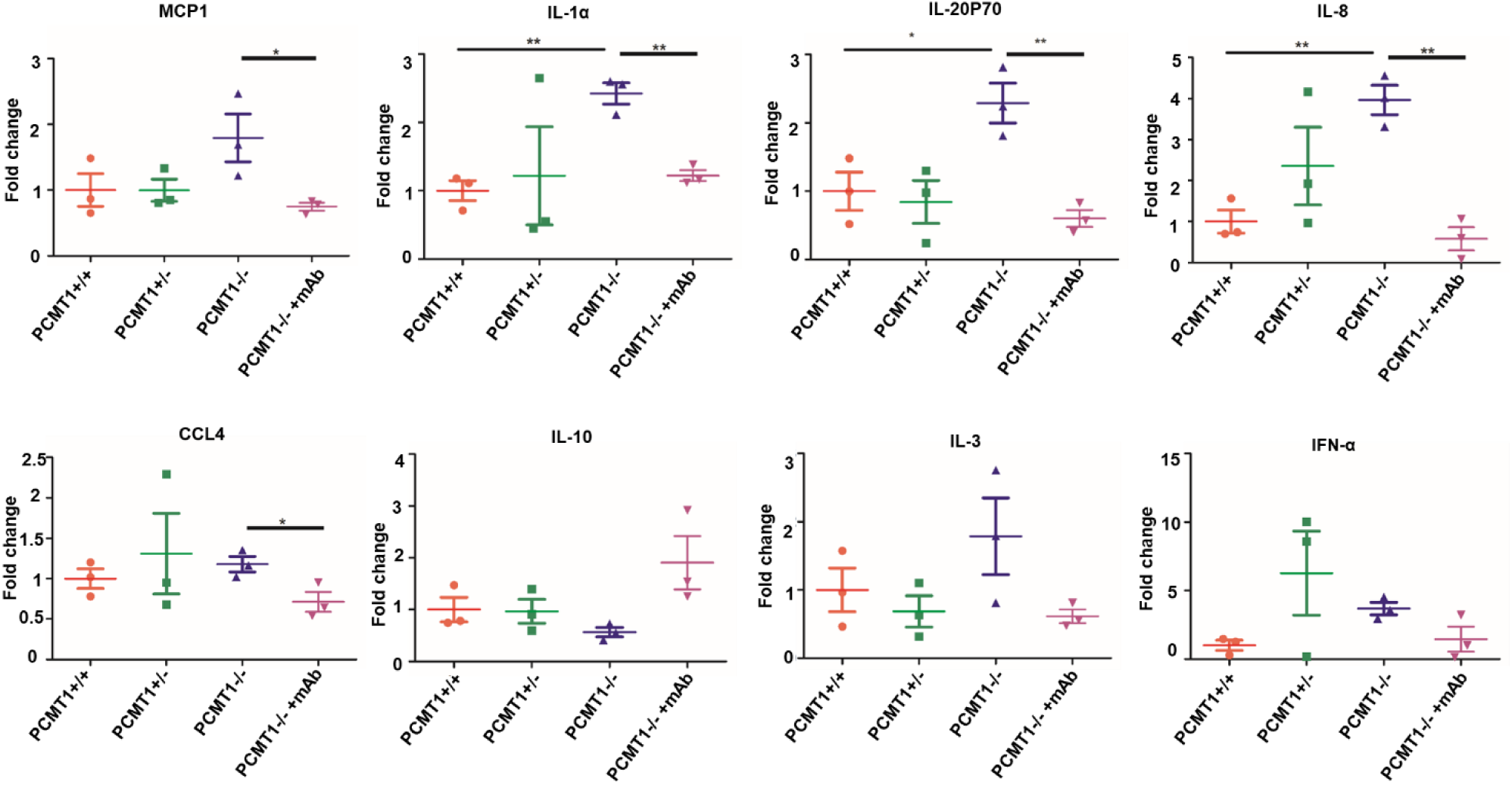
Anti-isoDGR mAb reduces levels of pro-inflammatory cytokines in lung of Pcmt1^-/-^ mice. Graph represents quantitative PCR analysis of pro- and anti-inflammatory cytokine expression in lung tissue from Pcmt1^+/+^, Pcmt1^+/-^, Pcmt1^-/-^, and mAb-treated Pcmt1^-/-^ mice aged 5-6 weeks. Expression of GAPDH was used to normalize data (n=3). Results are mean values ± SEM (* p<0.05, ** p<0.01, *** p<0.001).

### Synthetic isoDGR-modified peptides induce pulmonary inflammation

To confirm that isoDGR-modified proteins underlie the lung damage observed in Pcmt1^-/-^ mice, we next sought to replicate this pathology in 6-week-old Pcmt1^+/+^ mice (C57BL/6 background) via intranasal injection of synthetic isoDGR-peptide. Mice were administered 100µg isoDGR-peptide in 50µl PBS (or PBS-only vehicle control) and 24h later interstitial fluid was collected from the lungs for determination of pro-inflammatory cytokine levels. We detected significantly higher levels of inflammatory cytokines TNFα and IL-6 in lung fluid from isoDGR peptide-treated mice relative to those that received only vehicle control (Fig. S9B). Further analysis by immunostaining also revealed that residual isoDGR-peptides were co-localized with CD68^+^ immune cells in the lungs of treated mice (Fig. S9C-E), and that the extent of CD68 infiltration was positively correlated with isoDGR accumulation (Fig. S9F). Together, these results clearly indicate that isoDGR exerts pro-inflammatory effects in murine lungs.

### Transcriptomic profiling reveals TLR pathway activation in Pcmt1-KO mouse lung

Persistent low-grade inflammation in elderly individuals contributes to risk of age-linked disorders including CVD, cancer, and chronic pulmonary disease^29^. Previous studies have shown that damage-associated molecular patterns (DAMPs) can be recognized via Toll-like receptors (TLRs) that cross-talk with IL-1 signalling pathways and activate pro-inflammatory gene expression^30,31^. To further assess the pathological role of isoDGR *in vivo*, we next used RNA-seq to profile the lung tissue transcriptome of Pcmt1^+/+^, Pcmt1*^-/-^* and mAb-treated Pcmt1*^-/-^* mice. We identified a total of 13,920 genes in Pcmt1^+/+^ mice and 14,094 genes in Pcmt1*^-/-^* mice (13,611 genes were shared in common, while 443 genes were differentially expressed between genotypes) *(Fig.7A)*. Of the genes differentially expressed in Pcmt1*^−/−^* mice, 111 (∼25%) could be restored to Pcmt1^+/+^ levels by administration of anti-isoDGR mAb *(Fig.7A-B)*. Gene ontology analysis revealed that these 111 genes were critically involved in transcription, translation, and various other biological roles crucial for cell structure, function, and survival *(Fig.7B)*. When subjected to cluster analysis, we identified 30 upregulated genes in Pcmt1*^-/-^* mice, of which 21 could be restored to normal levels by mAb treatment. Similarly, of the genes downregulated in Pcmt1*^-/-^* mice, a majority of these were successfully normalised by mAb administration (12 of 21 genes; *Fig.7C-D)*. These transcriptomic data were then validated by RT-qPCR, which confirmed increased expression of FN1 and decreased levels of MT2, RGcc and PDK4 in the lungs of untreated Pcmt1^-/-^ mice (Fig. 7E-H). When we focused our analyses on TLR-mediated inflammatory pathways, we observed dysregulation of 31 key genes (21 increased, 10 decreased) in lung tissue from Pcmt1*^-/-^* mice compared to Pcmt1*^+/+^* mice. Similar to our previous analysis, 18 of 21 genes upregulated in Pcmt1^-/-^ mice were normalised upon mAb treatment (including potent effects on *TLR1, TLR2, TLR4, TLR6, Myd88* and *IRK4*), whereas 9 of 10 downregulated genes were restored to Pcmt1^+/+^levels by isoDGR-specific immunotherapy.

**Figure 7:**
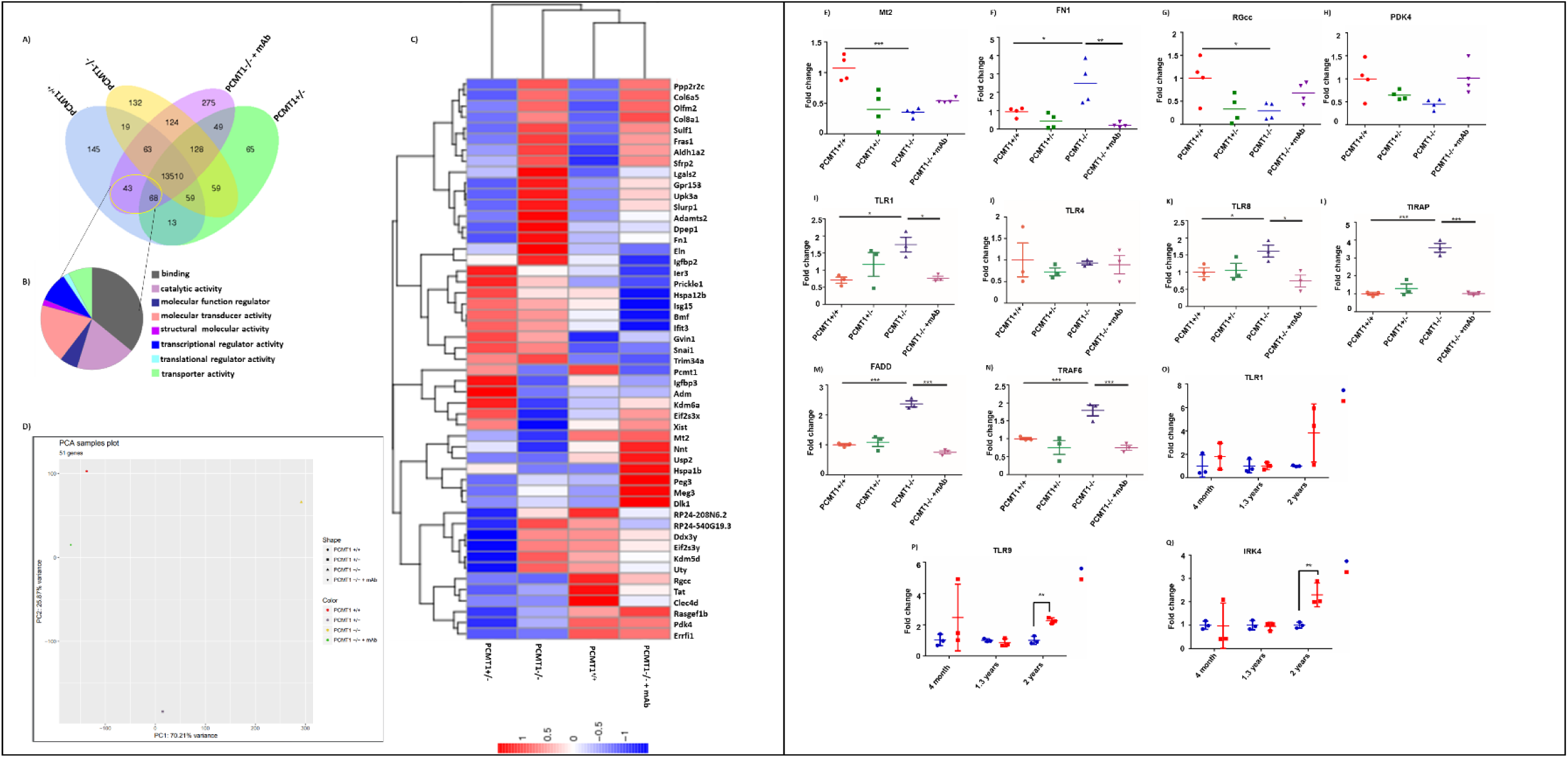
Lung transcriptome of Pcmt1^+/+^, Pcmt1^+/-^ Pcmt1^-/-,^ and mAb-treated Pcmt1^-/-^ mice. **(A)** Venn diagram indicating the number of differentially expressed genes across four samples (Pcmt1^+/+^, Pcmt1^+/-^, Pcmt1^-/-^ and mAb-treated Pcmt1^-/-^ mice) as well as overlap between each set of genes**. B)** Pie chart showing functional classification of 111 genes that were restored to normal expression levels in lungs from Pcmt1^-/-^ mice that received mAb treatment. **(C)** Hierarchical clustering of differential gene expression across all mouse lung samples in this study. Lung samples are represented by columns in the clustering heat-map and then sorted by genotype and treatment. **(D)** PCA plot for the 4 genotypes / treatment group. Each dot represents one group, color-coded according to genotype / treatment (Red: Pcmt1^+/+^; purple: Pcmt1^+/-^; yellow: Pcmt1^-/-^; Green: mAb-treated Pcmt1^-/-^). Graph shows quantitative PCR analysis of TLR signalling-associated genes TLR1 **(E)**, TLR4 **(F)**, TLR8 **(G)**, TIRAP **(H),** FADD **(I)** and TRAF6 **(J)** in lung from the four groups of mice at age 5-6 weeks. **(O-P)** TLR1, TLR9 and IRK4 relative expression in the lungs of Pcmt1^+/+^ and Pcmt^+/-^ mice at 4, 15, and 24 months old. Expression of GAPDH was used to normalize data. Results are mean ± SEM (* p<0.05, ** p<0.01, *** p<0.001).

### Aging-damaged isoDGR proteins activate TLR pathways

To confirm the role of isoDGR in activation of TLR pathways, we next performed RT-qPCR on total pulmonary RNA, which revealed increased expression of TLR1, TLR4, TLR8, TIRAP, FADD and TRAF6 in the lungs of Pcmt1^-/-^ mice (Fig. 7I-N). Strikingly, motif-specific mAb treatment was able to significantly reduce mRNA levels of these same TLRs in lung tissue from Pcmt1^-/-^ mice. Next, we studied isoDGR effects on TLR pathways using endothelial cells incubated on plates coated with native fibronectin (FN) or modified FN (isoDGR-FN) in the presence or absence of motif-specific mAb (or PBS-only control). RT-qPCR analysis revealed that HUVECs upregulated expression of TLR3, TLR4, TLR8 and ICAM when exposed to isoDGR- FN but not native FN or PBS only (Fig. S10). Addition of isoDGR-mAb efficiently blocked TLR induction in HUVECs co-cultured with isoDGR-FN. Consistent with these data, Pcmt1^-/-^ knock- down in HUVECs (HUVECPcmt1^-/-^) confirmed the ability of isoDGR to trigger TLR pathway activity, which was further associated with a significant reduction in proliferation and migratory activity relative to Pcmt1^+/+^ cells (Fig. S11). We then proceeded to quantify mRNA expression levels of TLR-associated genes in Pcmt1^+/+^ and Pcmt1^+/-^ mice at age 4, 15, and 24 months. These data revealed that TLR1, TLR9 and IRK4 expression levels increased over time in Pcmt1^+/-^ mice (Fig. 7O-Q). These results are in-line with previous reports that isoDGR-modified fibronectin can activate macrophages via integrin ‘outside-in’ signalling, thereby triggering an ERK:AP-1 cascade and pro-inflammatory release of MCP1 and TNFα^20^, which likely impact on TLR pathway activity^30–32^.

### IsoDGR-specific immunotherapy reduces Pcmt1^-/-^ lung pathology as determined by single-cell RNA sequencing

To gain a deeper understanding of isoDGR-induced pulmonary disorder in Pcmt1^-/-^ mice, we next conducted single-cell RNA sequencing of lung tissues from Pcmt1^+/+^, Pcmt1^-/-^, and mAb-treated Pcmt1^-/-^ mice at 6-weeks-old. We identified 29 distinct cell clusters, including lung-specific epithelial and structural cells, vascular and endothelial cells, as well as various leukocyte subsets (Fig. 8A). Among these clusters, B cells, T cells, and natural killer cells were markedly decreased in lung tissues from Pcmt1^-/-^ mice, whereas monocytes and dendritic cells were significantly increased, perhaps reflecting active infiltration by phagocytic cells and subsequent tissue inflammation. Imbalance of myeloid and lymphoid populations in the lung may be sufficient to enhance tissue injury and infection risk, while also inhibiting repair and resolution of inflammation^33^. Strikingly, mAb treatment restored both myeloid and lymphoid cell frequencies to the levels observed in Pcmt1^+/+^ mice (Fig. 8B). Our scRNA-seq data also revealed that Pcmt1^-/-^ mouse lungs exhibit increased numbers of pericytes, which are specialized cells that surround capillaries and regulate blood flow to downstream tissues^34,35^. High frequencies of lung pericytes in Pcmt1^-/-^ mice might indicate sub-optimal pulmonary circulation and endothelial barrier function, potentially related to the compromised gas exchange and increased lung permeability observed previously (Fig. 2). Together, these findings confirm that isoDGR accumulation in Pcmt1^-/-^ mouse lungs is associated with several pathophysiological defects, including blood vessel barrier issues, impaired gas exchange, and an altered immune profile that can be corrected by mAb treatment.

**Figure 8.**
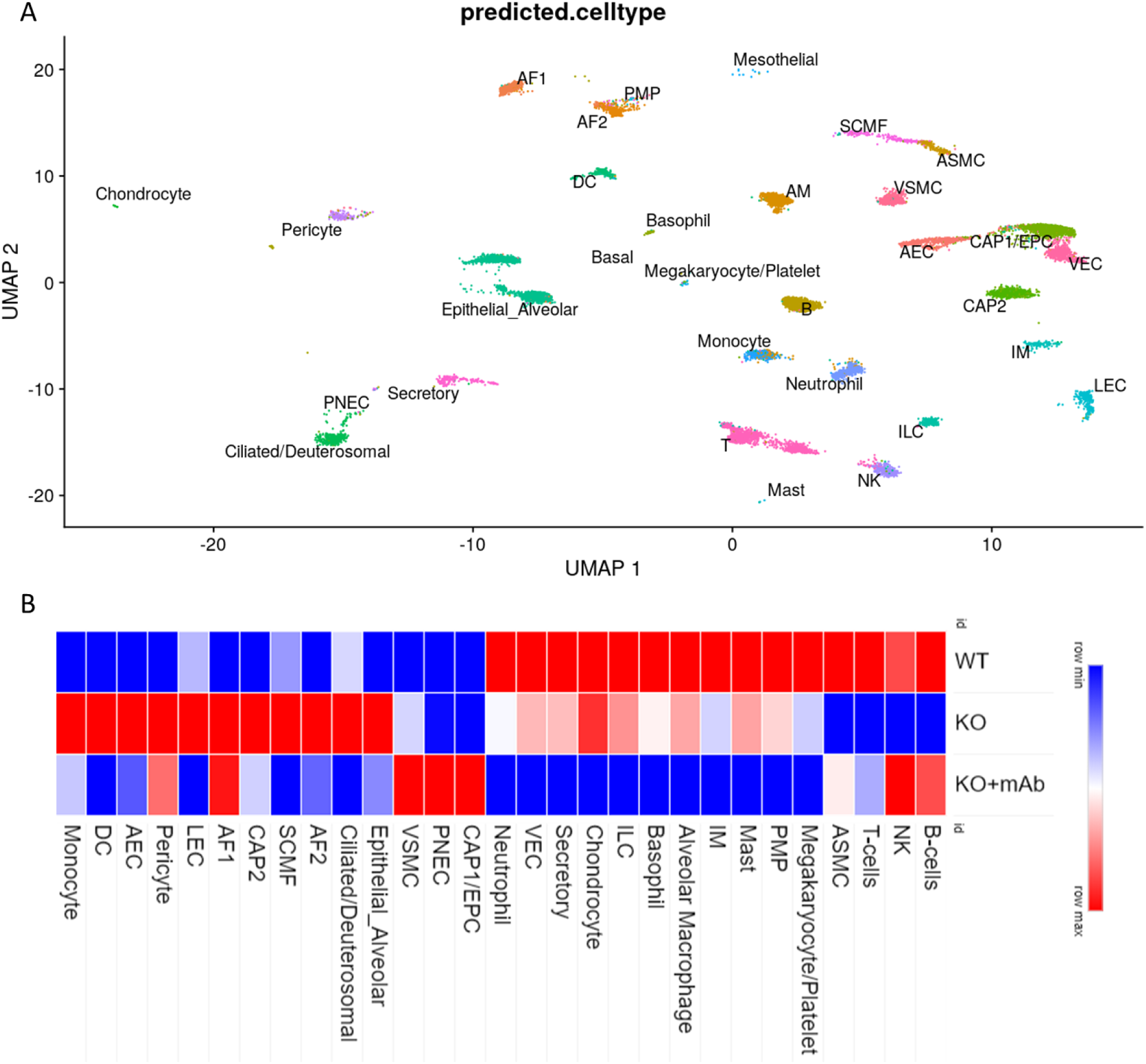
Lung cell landscape of Pcmt1^+/+^, Pcmt1^-/-^ and mAb-treated Pcmt1^-/-^ mice. (A) UMAP representation of the lung scRNA-seq results. Cell clusters were annotated with LungMap defined cell types using the Azimuth algorithm. A total of 29 different lineages were identified in murine lung. (B) Heat-map showing clusters and relative abundance of 29 lung cell types in the WT, Pcmt1 KO, mAb-treated Pcmt1 KO mice.

### Anti-isoDGR mAb treatment restores ribosomal protein expression in Pcmt1^-/-^ mouse lung

Pcmt1^-/-^ mice displayed significantly reduced weight compared to their control littermates. Among the genes found to be dysregulated upon scRNA-seq analysis of Pcmt1^-/-^ mouse lung we identified a subset of proteins associated with ribosomal machinery that were significantly decreased relative to littermate controls (including RPS35, RPS36, RPS38, RPS21, and RPS28; Fig. S12). Administration of anti-isoDGR mAb treatment appeared to increase expression of these ribosomal genes, so we next sought to validate the scRNA-seq data by RT-qPCR analysis of mouse lung tissues. This analysis confirmed that ribosomal gene expression was decreased in Pcmt1^-/-^ mouse lungs and could be partially restored by mAb treatment. Further studies will be necessary to fully elucidate how isoDGR-modified proteins impact the ribosomal machinery *in vivo*, since this finding may be directly linked with the reduced body weight and short lifespan of Pcmt1^-/-^ mice. Nonetheless, mAb-induced clearance of isoDGR-modified lung proteins was associated with marked weight gain and lifespan extension in Pcmt1^-/-^ mice, perhaps in-part due to restoration of protein synthetic functions.

### IsoDGR impairs pulmonary ETC and mitochondrial functions

Mitochondrial dysfunction is a hallmark of ageing and has been observed in both acute and chronic lung diseases^36^. Several reports have shown that ageing modulates mitochondrial DNA structure and functions, leading to dysregulation of energy metabolism (reduced ETC capacity and decreased ATP production) as well as enhanced ROS production^37–39^. Our RNA sequencing data indicated that ETC-linked gene expression was substantially reduced in the lungs of Pcmt1^-/-^ compared to control mice (Fig.9A). Validation of the RNA sequencing results by RT-qPCR also confirmed that lung tissue from Pcmt1^-/-^ mice displayed reduced expression of numerous ETC-associated genes (including ND1, ND2, ND3, ND4, NDL4, ND5, ND6, ATP6, Col1, Col2 and Cytb). Crucially, isoDGR-mAb administration to Pcmt1^-/-^ mice was able to restore normal expression levels of these dysregulated mtDNA genes, suggesting re-establishment of lung mitochondrial functions following treatment (Fig.9B). Since ETC defects are known to result in electron leakage, ROS generation, and inefficient respiration, we next performed seahorse and CellROX assays to determine whether isoDGR effects on mitochondria also impact energy metabolism. To do this, we seeded human lung endothelial cells (HULEC-5a) into PBS only, native FN, or isoDGR-modified FN, in the presence or absence of anti-isoDGR mAb. These experiments revealed that HULEC-5a cells exposed to isoDGR-FN displayed reduced oxygen consumption rate (OCR) and extracellular acidification rate (ECAR) relative to control cells (PBS only). However, isoDGR-specific mAb treatment was able to restore OCR and ECAR to normal levels in HULEC-5a cells (Fig.9C-D). Similarly, HULEC-5a cells generated significantly higher levels of ROS when cultured in the presence of isoDGR-FN, but this was significantly reduced by co-incubation with isoDGR-specific mAb (Fig. S13). Several reports have shown that ECM stiffness can trigger abnormal integrin signalling and mitochondrial dysfunction, thereby contributing to the development of disorders such as COPD, lung fibrosis, and emphysema^11,40–42^. These results are consistent with our present finding that isoDGR-modified ECM proteins promote age-linked mitochondrial dysfunction in Pcmt1^-/-^ mouse lung, whereas targeted antibody therapy can induce immune clearance of this damage motif and restore normal metabolic activity.

**Figure 9:**
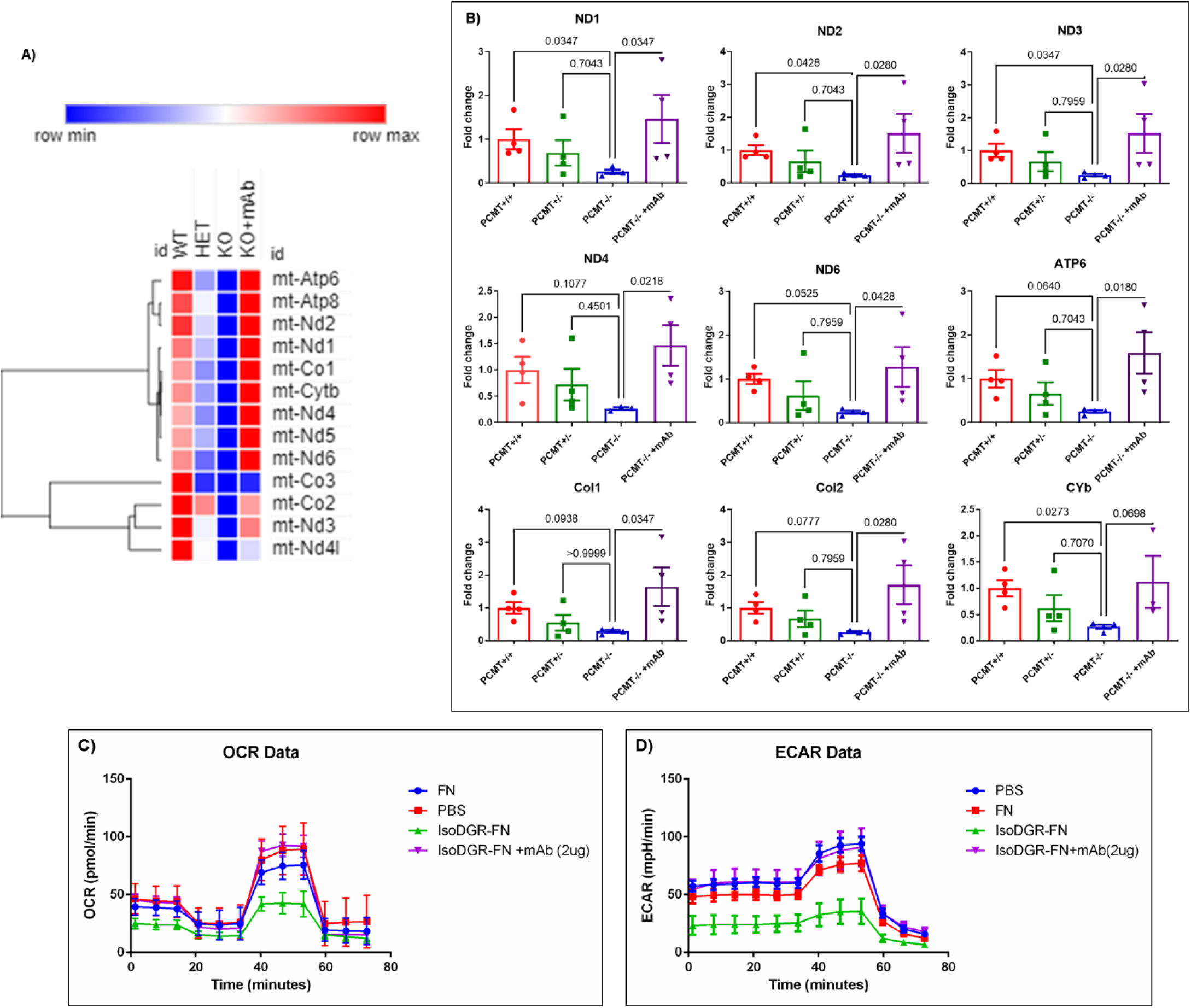
IsoDGR impairs ETC and mitochondrial functions in murine lung. **(A)** Heat-map of transcriptomic data showing that substantial down-regulation of mtDNA expression in lung tissue from Pcmt1^-/-^mice can be restored by anti-isoDGR mAb treatment. **(B)** Graph showing quantitative PCR analysis of mtDNA and ETC-related gene expression in the lungs of Pcmt1^+/+^, Pcmt^+/-^, Pcmt^-/-^, and mAb-treated Pcmt^-/-^ mice aged 6 weeks. Data were normalised to GAPDH expression level. Statistical significance was determined by Kruskal-Wallis test. Results shown are mean values ± SE. **(n=4)(C & D)** HULEC-5a were seeded into Seahorse Bioscience microplates for assessment of mitochondrial function-associated OCR **(C)** and ECAR **(D)** using a XF96 extracellular flux analyser (n=12- 16).

**Figure 10:**
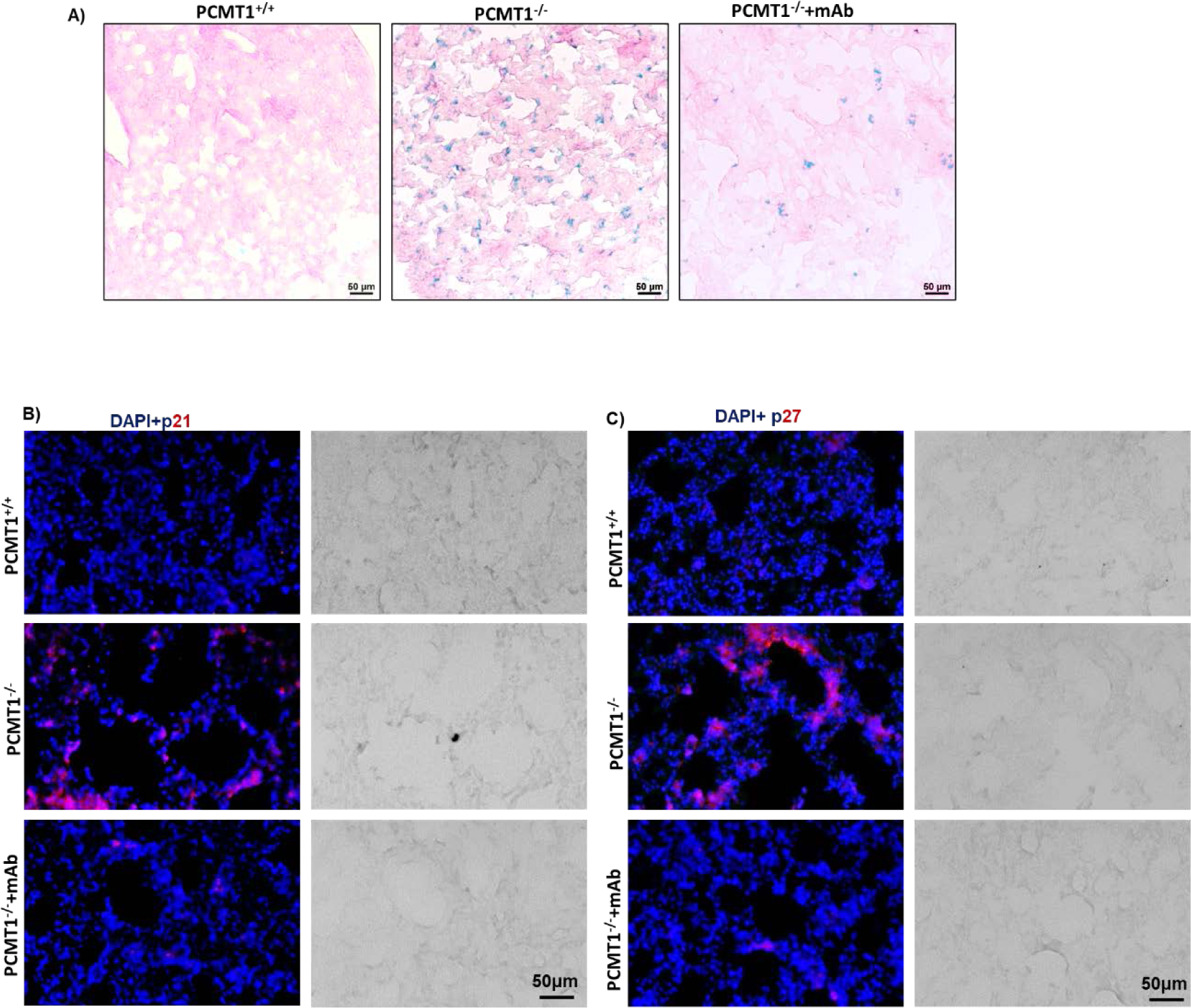
IsoDGR promotes senescence of lung parenchymal and EC cells in Pcmt1^-/-^ mice. Representative immunostaining images showing β-gal **(A)**, P21 **(B)** and P27 **(C)** levels in lung tissue from the respective genotypes / treatment conditions (n=5)

### IsoDGR promotes senescence and apoptosis of lung parenchymal and endothelial cells

ROS induced by smoking, environmental, and other oxidative stresses are known to drive inflammation and cellular senescence in lung tissues, leading to chronic lung pathologies^36,43,44^. To test whether ROS arising from isoDGR accumulation can also promote cellular senescence and apoptosis, we performed β-galactosidase (β-gal) and Tunel staining of lung tissue from Pcmt1^+/+^, Pcmt1^-/-^, and mAb-treated Pcmt1^-/-^ mice. We observed significantly increased β-gal levels (Fig.14) and apoptotic cell numbers (Fig. S14) in lung parenchyma and endothelium from Pcmt1^-/-^ mice, whereas mAb treatment substantially reduced these pathological features. Consistent with the β- gal and Tunel staining, we also observed increased expression of P21, P27 and caspase-3 in lung tissue from Pcmt1^-/-^ mice (Fig 11 & S14), as well as elevated frequencies of apoptotic lung parenchymal and endothelial cells in 2-year-old Pcmt1^+/+^ and Pcmt1^+/-^ mice relative to young mice (Fig. S14c). These data further indicated that relative levels of isoDGR accumulation over the course of natural aging directly correlate with cellular dysfunction and tissue degradation *in vivo*.

Indeed, our results are consistent with reports that protease-mediated clearance of age-damaged proteins can reduce cellular senescence and apoptosis to improve tissue function^45^. Together, these findings clearly indicate that pulmonary accumulation of isoDGR-modified proteins leads to increased cellular senescence and apoptosis, whereas motif-specific mAb can promote immune clearance of damaged proteins and reduce lung pathology.

## Discussion

Aging is the primary risk factor for many chronic diseases that increase risk of early mortality. As the global population becomes more elderly, current projections suggest this will drive an exponential increase in worldwide burden of age-linked diseases and associated healthcare costs^46,47^. Consequently, there is an urgent need to better understand the complex processes that underpin human aging, so that we can begin to develop methods of extending health lifespan in elderly cohorts. Current models suggest that time-dependent accumulation of molecular damage leads to degradation of tissue function as humans age^48^. In particular, age-linked degenerative protein modifications (DPMs) have been shown to alter protein structure and bioactivity *in vivo*, leading to increased risk of several major diseases^48,49^. While selective treatment can reduce symptoms of specific disorders, targeting shared molecular mediators of multiple age-linked pathologies may be more effective for extending human ‘health-span’.

The ECM is a non-cellular component of all tissues and organs that critically supports many key biochemical and biomechanical processes. ECM proteins exhibit slower turnover than cellular proteins throughout the human body, and are thus more susceptible to accumulation of age-associated DPMs. While the effects of age-related molecular damage can be seen in all organs, some tissues may be more seriously impacted by DPMs than others. In particular, ECM-rich organs including the skin, lung, and blood vessels are likely to be highly susceptible to DPM accumulation, leading to time-dependent decline in critical functions including barrier maintenance, gas exchange, and delivery of oxygen / nutrients to distant body tissues.

We previously reported that fibronectin, laminin, tenascin C proteins in carotid atherosclerotic plaque from CVD patients exhibit marked deamidation of NGR sequences, thereby forming integrin-binding isoDGR motifs that facilitate increased leukocyte adhesion^19,20^. These age-damaged matrix proteins enhance binding interactions between monocytes, macrophages, and endothelial cells, while also triggering ‘outside-in’ integrin signalling to elicit a range of potent cytokines and chemokines^19,20^. The lung ECM is essential for normal organ function and comprises a complex mixture of fibrous proteins (collagen, elastin), glycoproteins (fibronectin, laminin), glycosaminoglycans (heparin, hyaluronic acid) and proteoglycans (perlecan, versican)^8^. Accordingly, ECM dysregulation is now recognised as a key feature of lung aging^8^, and we have previously detected isoDGR-modified ECM proteins in blood vessels^20^, which form an integral part of pulmonary anatomy. Consistent with these data, the current study demonstrates that age-linked accumulation of isoDGR can trigger a range of pathological features of major lung diseases such as fibrosis, asthma, emphysema, COPD, and cancer^50^ (including pulmonary inflammation, oedema, hypoxemia, congestion of blood vessels, coagulopathy, mitochondrial dysfunction, oxidative stress, and cellular senescence). Intriguingly, while isoDGR accumulates in all body tissues of Pcmt1^-/-^ mice, the severity of motif-induced lung pathology is strongly correlated with overall mortality, indicating that pulmonary dysfunction is a key determinant of lifespan in this model. Crucially, our results indicate that motif-specific mAb therapy was able to induce immune clearance of age-linked isoDGR damage and increased longevity of treated mice^12^. This immunotherapeutic strategy could potentially also be used in clinical settings to help maintain health and reduce age-linked disease burden in elderly cohorts.

Biomolecular damage (including DPM accumulation) has long been recognized as a key mediator of human aging and chronic disease. However, the mechanisms by which DPMs induce age-linked pathology and the primary organs impacted by this process have remain elusive. Since DPMs are not encoded by the genome or generated by enzymes, they cannot be studied using common genetic approaches. Consequently, progress in this area has been severely restricted by the technical challenges of identifying key proteins and modification sites against a complex backdrop of widespread protein damage *in vivo*. However, advanced LC-MS/MS proteomics now enables the detection of many DPMs in human clinical samples, and new bioinformatic tools can assist in identifying DPMs with likely functional impact on human health and disease. In the current study, we investigated isoDGR accumulation in naturally-aged mouse and human tissues, then uncovered a range of novel motif functions combining the Pcmt1^-/-^ mouse model with target-specific monoclonal antibodies for biochemical / immunological assays *in vitro*. In addition, we further demonstrated that a motif-specific antibody can also be used to treat isoDGR-induced disease by promoting phagocytic clearance of age-damaged proteins from the lungs *in vivo*. The strategies developed in this study open new avenues for future investigations of DPM biology in health and disease, and could potentially be used to develop new interventions in age-linked human disorders.

## Materials and methods

### Human lung tissue microarrays

We obtained two types of paraffin-embedded human lung tissue array (TMA-LC2086a and TMA- LC561) from Tissue Array UAS. TMA-LC2086a contains of 192 sections of normal human lung tissues with varying age, whereas TMA-LC561 consists of 56 sections of pulmonary interstitial fibrosis tissues, alongside 2 cases each of cancer-adjacent lung tissues and normal lung tissues. Our objective was to analyse isoDGR-modified protein accumulation in aging human lung tissues and assess potential association with CD68+ and CD11b+ cell distribution. To prepare the TMA slides for immunostaining, they were first deparaffinized and rehydrated by sequential immersion in 100% ethanol, followed by 75%, 50% ethanol, and finally PBS. Heat antigen retrieval was performed by placing the slides in a Pyrex beaker with antigen retrieval solution, and the temperature was raised to 95-99°C for 30-60 minutes. The beaker was covered with cling wrap to maintain temperature, and the slides were then removed and placed into a fresh coplin jar. After washing with milli Q H2O for 5 minutes, the slides were washed with PBS-TT (0.5% Tween-20, 0.1% Triton X-100) for 15 minutes. Next, the slides were incubated with a protein block (2.5% normal goat serum, 1% BSA in 0.5% PBST) at room temperature for 20 minutes. After drying the slides, DAPI staining was applied for 15 minutes to determine tissue autofluorescence, which was later used to subtract tissue background from the antibody-stained images. For immunostaining, the slides were treated with AlexaFluor-conjugated primary antibodies (isoDGR-AlexaFluor 488 [1:200], CD68-AlexaFluor 595 [1:200], and CD11b-AlexaFluor 700) overnight at 4°C. After immunostaining, the slides were washed 3 times using 1X PBS and mounted with aqueous mounting media. Finally, images were acquired using an Axion scan.Z1 slide scanner for further analysis.

### Animals

WT C57BL/6J mice at 17 months of age (catalog number: 000664), and Pcmt1^+/−^ (C57BL/6 background) mice (B6;129S4-Pcmt1tm1Scl/J; Strain #:023343; RRID:IMSR_JAX:023343) were obtained from the Jackson Laboratory in Bar Harbor, ME, USA. Animals were housed under specific pathogen-free (SPF) conditions in isolator cages for two weeks prior to the start of the experiments. The animal house maintained at room temperature of 24°C with 12 hour light/dark schedule. Mice were provided access to Purina rodent chow diet and tap water *ad libitum*. Mice deficient in deamidation repair enzyme Pcmt1 were previously generated by Clarke and co-worker^25^. The Pcmt1^+/−^ (male and female) mice were bred to yield litters comprising Pcmt1^+/+^, Pcmt1^+/−^, and Pcmt1^−/−^ offspring in the expected Mendelian ratios. Mouse genotype was confirmed by PCR using the primers as shown in Table S1. All animal experiments used age- and sex-matched mice unless otherwise specified. All mouse procedures were performed in a humane manner and approved by the Nanyang Technological University Institutional Animal Care and Use Committee (IACUC protocol # ARF-SBS/NIE-A18016, ARF-SBS/NIE-A19029) or the Animal Care Committee at Brock University (AUP # 22-08-04).

### Lung tissue H&E and immunostaining

Lungs from Pcmt1^+/+^, Pcmt1^+/-^, Pcmt1^-/-^, and mAb-treated Pcmt1^-/-^ mice were collected and fixed with 4% PFA at 4°C for 24h. Tissues were washed with 1X PBS and transferred into 15% sucrose, then 30% sucrose, and kept at 4°C. Lungs were subsequently embedded in OCT compound with dry ice and sectioned (10µm) using a Leica CM3060S Cryostat. Sections were mounted onto Fisherbrand Superfrost Plus microscope slides, then kept in warm PBS for 20min to remove OCT prior to staining with haematoxylin and eosin. For immunostaining, slides were permeabilized with 0.5% PBST for 2-3h and then incubated with blocking buffer (2.5% normal goat serum, 1% BSA in 0.5% PBST) for 1h at RT prior to addition of primary antibodies (isoDGR [1:200] and CD68 [1:200]) overnight at 4°C. Slides were washed with PBS (3X) for 5min, then incubated with secondary antibodies conjugated to AlexaFluor 488 and 594 (1:500) for 1h at RT. Immunostained slides were washed with PBS (3X) for 5min then incubated with DAPI for 15min to visualize cell nuclei. After staining, slides were washed with 1X PBS and mounted with aqueous mounting media. Images were acquired using a Zeiss LSM710 confocal microscope.

### MRI analysis

Mouse lungs of different genotypes from the same litter were excised for MRI imaging. All MRI experiments were conducted on a 14 Tesla Bruker Ascend 600WB vertical magnet, equipped with a MicWB40 micro-imaging probe and a Micro2.5 gradient system. A quadrature coil with inner diameter of 30mm (MICWB40 RES 600 1H 040/030) was used to transmit/receive MR signals. Images were obtained using Paravision v6.0.1 software. A modified multi-spin-multi-echo (MSME) pulse sequence was used with an echo time (TE) of 8ms, a repetition time (TR) of 3000ms, with 2 averages to acquire images from the excised mouse lungs (positioned in a 30mm diameter glass tube). The tissue was surrounded by fomblin® to suppress background signals and placed horizontally such that axial images provided a cross-section of both lobes. Each slice thickness was 0.4mm, with a 22 x 22mm field of view, and each scan was captured as a 512 x 512 pixelated image.

### Western blot analysis

Western blot analysis was performed using standard methods. Primary antibodies and dilutions were as follows: isoDGR (mouse monoclonal 1:1000), Pcmt1 (rabbit polyclonal, Abcam 1:1000), GAPDH (Invitrogen 1:1000).

### SpO2 assay

We used a rodent oximeter to measure SpO2 in Pcmt1^+/+^, Pcmt*1^+/-^*, Pcmt1^-/-^ and mAb-treated Pcmt^-/-^ mice according to manufacturer’s protocol.

### Evans blue lung permeability assay

Mice were anaesthetized by isoflurane inhalation and 200µl 0.5% Evans blue in 1X PBS was administered via retro-orbital route. After 30min, mice were perfused with 15ml of 1X PBS. Organs were collected and air dried to eliminate water content variability between different tissues. A 500µl volume of formamide was added to 50-100mg lung tissue and incubated at 55°C for 24h to extract Evans blue. After removing tissue fragments from the sample tube, absorbance was measured at 610nm using neat formamide as a blank. The final amount of Evans blue extravasation was calculated as ng dye per mg of tissue. The experiment was performed with n=3 animals.

### Collection of bronchoalveolar lavage (BAL) fluid for cytokine assay

Synthetic isoDGR-peptide (100µg) in 50µl of PBS (or PBS only vehicle control) were intranasally injected into the lungs of Pcmt1^+/+^ mice (C57BL/6 background). After 24h, mice were anaesthetized by ketamine / xylazine injection and 200µl PBS was used to lavage the lungs three times. Lavage fluid was then centrifuged and the cell-free supernatants collected for cytokine analysis.

### Cytokine multiplex bead assay

The LEGENDplex^Tm^ mouse inflammation panel (Biolegend, San Diego, CA) was used to measure 13 different cytokines (IL-23, IL-1α, IL-1β, IL-6, IL-10, IL-12p70, IL-17A, IL-23, IL-27, MCP1, IFN-β, IFN-γ, TNF-α, and GM-CSF) in plasma from Pcmt1^+/+^, Pcmt1^+/-^, Pcmt1^-/-^ and mAb-treated Pcmt^-/-^ mice (assessed by LSRII flow cytometer according to the manufacturer’s protocol).

### Terminal deoxynucleotidyl transferase dUTP nick end labeling (TUNEL) assay

Lung sections were immersed in PBS for 5min (2X) and incubated with proteinase K (1:100 in 10mM Tris pH 8) for 20min at room temperature. Slides were rinsed with PBS then incubated with equilibration buffer for 20min. Sections were next incubated with labeling reaction mixture (equilibration buffer 40µl, terminal deoxyribonucleotidyl transferase [TdT] enzyme buffer 5µl, labeling solution 5µl) at 37°C for 1h. Slides were then washed with PBS (3X) for 5min prior to incubation with DAPI for 10min followed by a further PBS wash for 5min. The slides were mounted with glycerol-based mounting media. TUNEL-positive apoptotic cells were observed and images were acquired using a Zeiss LSM710 confocal microscope.

### Senescence-associated β-galactosidase (SA-βgal)

Cryosections of frozen lung tissue were fixed with 1% PFA for 10min and washed in PBS (3x at 5 min/wash) then incubated overnight at 37°C with staining buffer (5-bromo-4-chloro-3-indolyl β-D-galactopyranoside [X-gal, Bioline, London, UK], 5mm K3Fe[CN]_6_, 5mm K4Fe[CN]_6_, and 2mm MgCl_2_ in PBS at pH 6.0). After washing with PBS, the sections were mounted with glycerol-based mounting media and imaged using bright field microscopy.

### Cell culture and generation of Pcmt1 knock-down cells by shRNA

HUVECs were cultured in DMEM containing 10% FBS and 1% penicillin-streptomycin at 37°C in a CO_2_ incubator. Pcmt1 shRNA in PLKO plasmid (GCGCTAGAACTTCTATTTGAT, TRCN0000036400, Sigma-Merck USA) was transfected into HUVECs. After 24h, Puromycin (2 µg/mL) was used to select stably transformed cells. Pcmt1 knock-down was confirmed by western blot.

### CellROX® ROS assay

A total 12,000 HULEC-5a cells were seeded into each chamber of a 96 well plate (each chamber was coated with native FN, isoDGR-FN, or PBS control in the presence or absence of 2ug/ml anti-isoDGR mAb. After 24h, 5µM CellROX red reagent (Thermo Fisher Scientific, Inc.) was added to the HULEC-5a cultures and incubated for 30min in a 5% CO_2_ incubator at 37°C. The cells were then washed with PBS, fixed with 3.7% formaldehyde in PBS for 15min, and nuclei stained with DAPI for 5min at room temperature. ROS positive cells were imaged using a Zeiss LSM710 confocal microscope.

### RNA-seq analysis

RNA-seq was performed by Novoseq Technology, Singapore. Four libraries of transcriptomic samples were generated from the lungs of Pcmt1*^+/+^*, Pcmt1*^+/−^*, Pcmt1*^−/−^*, and mAb-treated Pcmt1*^−/−^*mice. The length of RNA fragments was detected using an Agilent 2100 Bioanalyzer (Agilent, USA). The cDNA library was then constructed using PCR amplification. RNA-seq was carried out with the Novogene strategy on an Illumina second-generation high-throughput sequencing platform. Reads with inferior quality or adapters were filtered. Clean reads data were processed using Tophat2 and Cufflinks software to complete the alignment of transcriptomes and transcript splicing analysis separately. Clean reads were mapped to the *Mus musculus* reference genome (GRCm38.p6/NCBI, GCF_000001635.26).

### Differentially expressed genes (DEG) analysis

Normalized fragments per kilobase of exon per million mapped reads (FPKM) was used to calculate gene expression level with the union counting model in HTSeq software (Princeton University, USA). We applied a cut-off value of FPKM > 1 to define gene expression, and DEG analysis was performed using DESeq (Bioconductor, USA). Genes with differential expression were screened and hierarchical clustering analysis was performed.

### Lung cell preparation and single cell RNA sequencing

Lung tissues were collected from 6-week-old Pcmt1^+/+^, Pcmt1^-/-^, and mAb-treated Pcmt1^-/-^ mice then transferred into RPMI-1640 medium containing 10% FBS and kept on ice. The tissues were washed with 1X PBS and then chopped into small pieces with scissors. Next, the tissue was resuspended in 1ml digestion buffer consisting of collagenase type I (2 mg/ml), hyaluronidase (1 mg/ml), and DNase I (0.2 mg/ml) in RPMI-1640 medium before incubation at 37°C for 30–40 minutes with gentle agitation at 110 RPM. The tissue suspension was then gently pipetted up and down, and the resulting cell suspension was passed through 100μm mesh filters. The cell suspension was quenched with 10 ml of RPMI containing 10% FCS. After centrifugation at 500 × g for 3 minutes at 4°C, the cell pellet was re-suspended in RPMI medium containing 10% FBS for scRNA-seq.

### Single cell RNA sequencing and data analysis

Cells were subjected to droplet-based single cell sequencing. The cDNA libraries were prepared using a Chromium Single Cell 3’ V3 kit (10X Genomics, USA) according to the manufacturer’s instructions and then sequenced on a NovaSeq6000 (Illumina, USA). Raw sequencing reads were aligned to the mm10 (GENCODE vM23/Ensembl 98) mouse reference genome using Cell Ranger (v7.1.0) to generate single-cell count matrices that were normalized, integrated, and annotated using Seurat (v4.30)^51^. Low-quality cells were filtered using uniform quality control thresholds; cells with RNA counts less than 300 and with mitochondrial read percentages more than 20% were filtered out. A total of 29653 cells were obtained across all 3 conditions. Log normalization was implemented in Seurat. Principal component analysis (PCA) was performed to reduce the dimensionality of each dataset, and the first 35 PCA components were used to construct UMAP. Unsupervised cell clustering was performed with the Louvain algorithm. Cluster marker detection was performed by differentially expressed gene (DEG) analysis for each marker against the remaining markers using FindAllMarkers. Azimuth scRNA-seq references for lung cells were downloaded from the LungMAP portal. Cell type annotation was performed automatically using LungMAP Cell Reference with Azimuth,^52^ since LungMap includes comprehensive annotation of endothelial cells, epithelial cells, mesenchymal cells and immune cells in lung tissue^53^.

### Reverse transcription-quantitative polymerase chain reaction (RT-qPCR)

Total RNA isolated from lung tissues was treated with DNase and reverse-transcribed using a first-strand DNA synthesis kit from Invitrogen. PCR was performed on an ABI Fast 7500 System (Applied Biosystems, Foster City, CA). TaqMan probes for the respective genes were custom-generated by Applied Biosystems based on sequences from the Illumina array and used as per the manufacturer’s instructions. Expression levels of target genes were determined in triplicate from the standard curve and normalized to GAPDH mRNA level. The RT-PCR primers are exhibited in Table 2 and 3.

### Seahorse assay for metabolic flux analysis and glycolysis stress test

The metabolic flux analyses and glycolysis stress test were performed using a Seahorse XF96 (Agilent Technologies, USA). In the metabolic flux analysis, oxygen consumption rate (OCR) was measured to assess mitochondrial function in HULEC-5a cells. In brief, 12,000 cells per well were seeded into MCDB media (Merck, Singapore) using an XF96 well plate (Agilent, Santa Clara, CA) coated with PBS control, native FN, or isoDGR-FN, in the presence or absence of 2ug/ml anti-isoDGR mAb. After 1h, the medium was switched to Seahorse XF DMEM (Agilent), containing 10ng/mL Epidermal Growth Factor (EGF), 1 µg/mL Hydrocortisone, and 10mM Glutamine, then incubated in a non-CO_2_ incubator. The Seahorse XF96 Analyzer (Agilent) was used to measure extracellular acidification (ECAR) and oxygen consumption rate (OCR) in real-time. The OCR assay consisted of 3 distinct injections. The first injection of 1μM oligomycin blocks mitochondrial ATP production; the second injection of 1.5μM FCCP (carbonyl cyanide-4 (trifluoromethoxy) phenylhydrazone) uncouples the mitochondrial membrane to assess maximal cellular respiration; and the third injection of 100μM rotenone together with 1μM antimycin A blocks ETC complexes I and III, respectively. For glycolysis stress test, a similar procedure was performed except that non-glycolytic acidification was measured first in the absence of both glucose and pyruvate. Final concentrations of 10mM glucose (Sigma Aldrich, St. Louis, MO), 1µM oligomycin (Sigma Aldrich), and 5mM 2-deoxy-D-glucose were added to assess ECAR.

### Statistical analysis

Statistical analyses were performed using GraphPad Prism v9.0 (GraphPad Software, Inc., San Diego, CA). Data were tested for normality using D’Agostino & Pearson, or Shapiro-Wilk tests. For normally distributed data, differences between groups were assessed either by two-tailed unpaired Student’s t-test or one-way ANOVA for multiple groups, followed by Tukey’s multiple comparisons test. Otherwise, Kruskal-Wallis test with Dunn’s multiple comparisons post-hoc test was used (*p*<0.05 was considered significant).

## Data Availability

RNA-seq data presented in this study are deposited in GEO-NCBI (GSE224434). All other data are included in the main text or supplemental materials.

## Acknowledgments

This work was supported in part by the Singapore National Medical Research Council (NMRC/OFIRG/0003/2016), Canadian Institutes of Health Research Tier1 Canada Research Chair (CRC-2020-00263), Canadian Institutes of Health Research Project Grant (PJT-186091), The Natural Sciences and Engineering Research Council of Canada Discovery Grant (RGPIN-2023-04304), Canada Foundation for Innovation Grant, Ontario Research Fund, and start-up research grant from Brock University.

## Author Contributions

K.P., N.M. and S.K.S. designed the project. K.P., K.M., S.C.N., R.I., B.S.L., B.S., R.M., S.T.B., performed the experiments and data analysis; P.K. and S.K.S. drafted the manuscript; E.S.C., V.A.F., R.E.K.M., J.L., P.K., E.L.T., K.L.L., I.H.S., Y.G.G., A.M.R, R.N.K. and C.C. provided resources/equipment; R.E.K.M., J.L., P.K., E.L.T., Y.G.G., K.L.L., N.M. and S.K.S contributed grants/reagents/materials/analysis tools. K.P., N.M. and S.K.S. conceived and supervised the project. All authors revised the manuscript.

