## Supplemental Figures and Tables for "Age-linked lung pathology is reduced by immunotherapeutic targeting of isoDGR protein damage"

**Supplemantry Data**

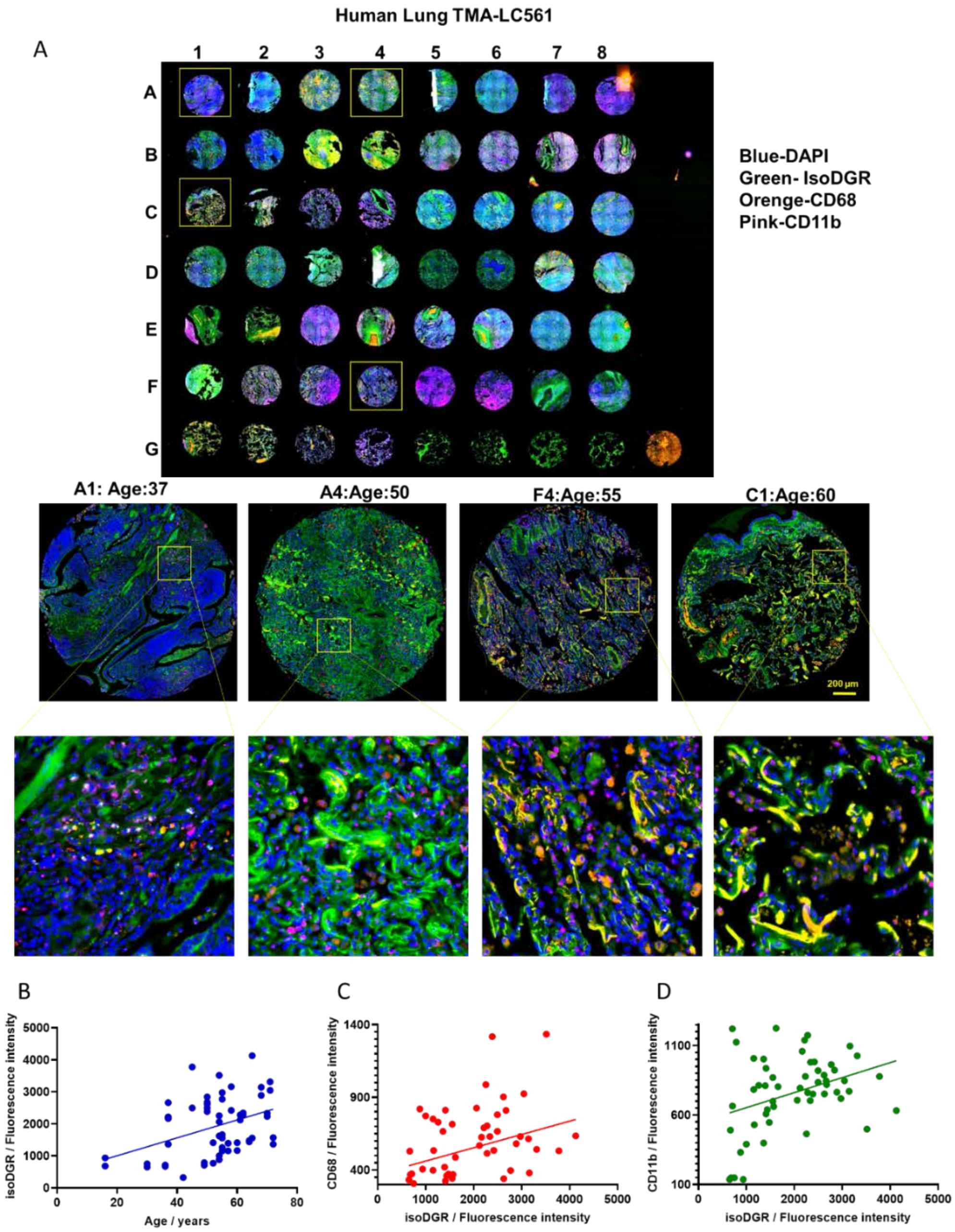


***Figure S1: Age-induced accumulation of isoDGR-modified proteins correlates with CD68+ and CD11b+ cells in human lung tissue.*** *(A) Representative immunostaining images showing isoDGR-protein distribution and correlation with CD68+ and CD11b+ immune cells in 56 sections of pulmonary interstitial fibrosis tissues, alongside 2 cases each of cancer-adjacent lung tissues and normal lung tissues of varying age. Representatives zoomed images are shown; A1: Age 37, A4: Age50, F4 Age 55, C1: Age 60. (B) IsoDGR level was positively correlated with age (linear regression slope 26.20). (C) CD68 level was positively correlated with isoDGR level. (D) CD11b level was positively correlated with isoDGR level.*

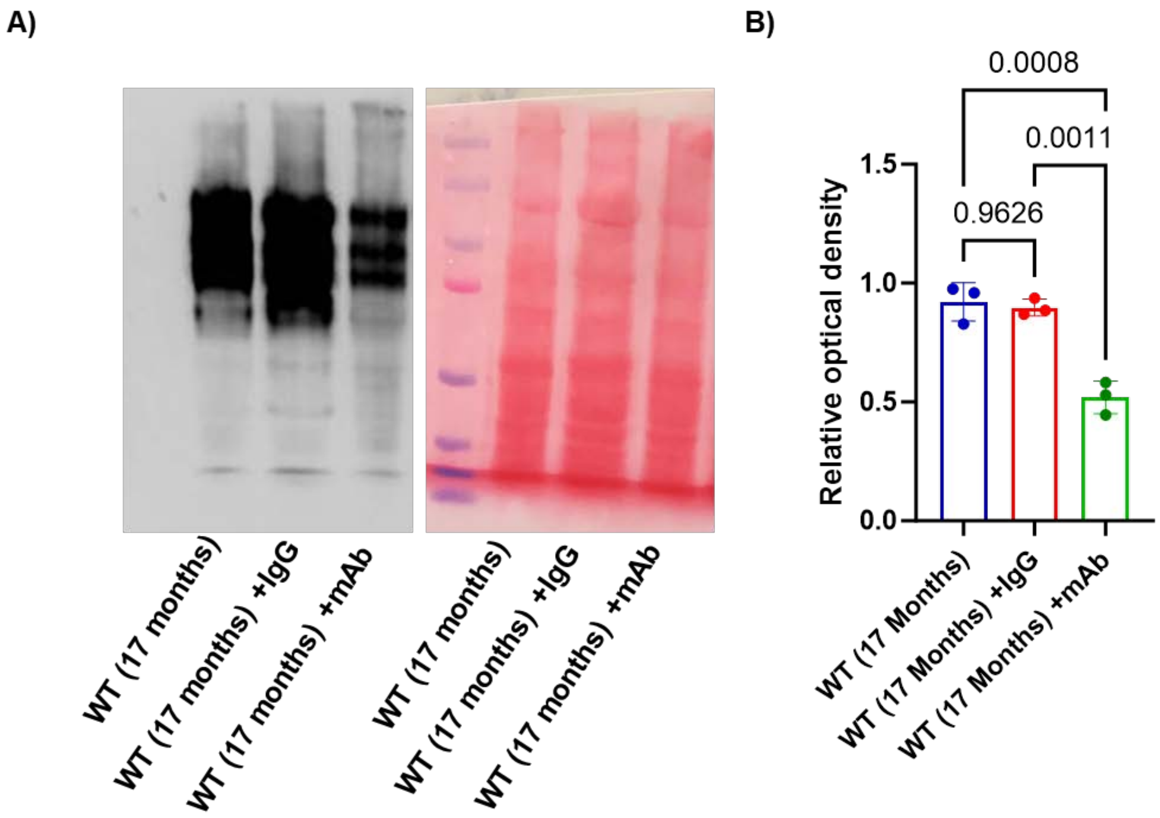


**Figure S2** ***Immune clearance of isoDGR-damaged proteins in lung tissues of naturally-aged mice***

***(A)*** *Lung protein lysates from 17 months old WT mice, 17 months old WT mice treated with isotype IgG antibody, and 17 months old WT mice treated with isoDGR-specific mAb subjected to western blot using isoDGR-specific mAb. Protein loading was visualized by Ponceau S.(n=5)* ***(B)*** *Graph showing quantification of isoDGR-damaged protein levels in the lungs of these mice (n=5) assessed at 17 months weeks. Results are shown as mean ± SEM.*

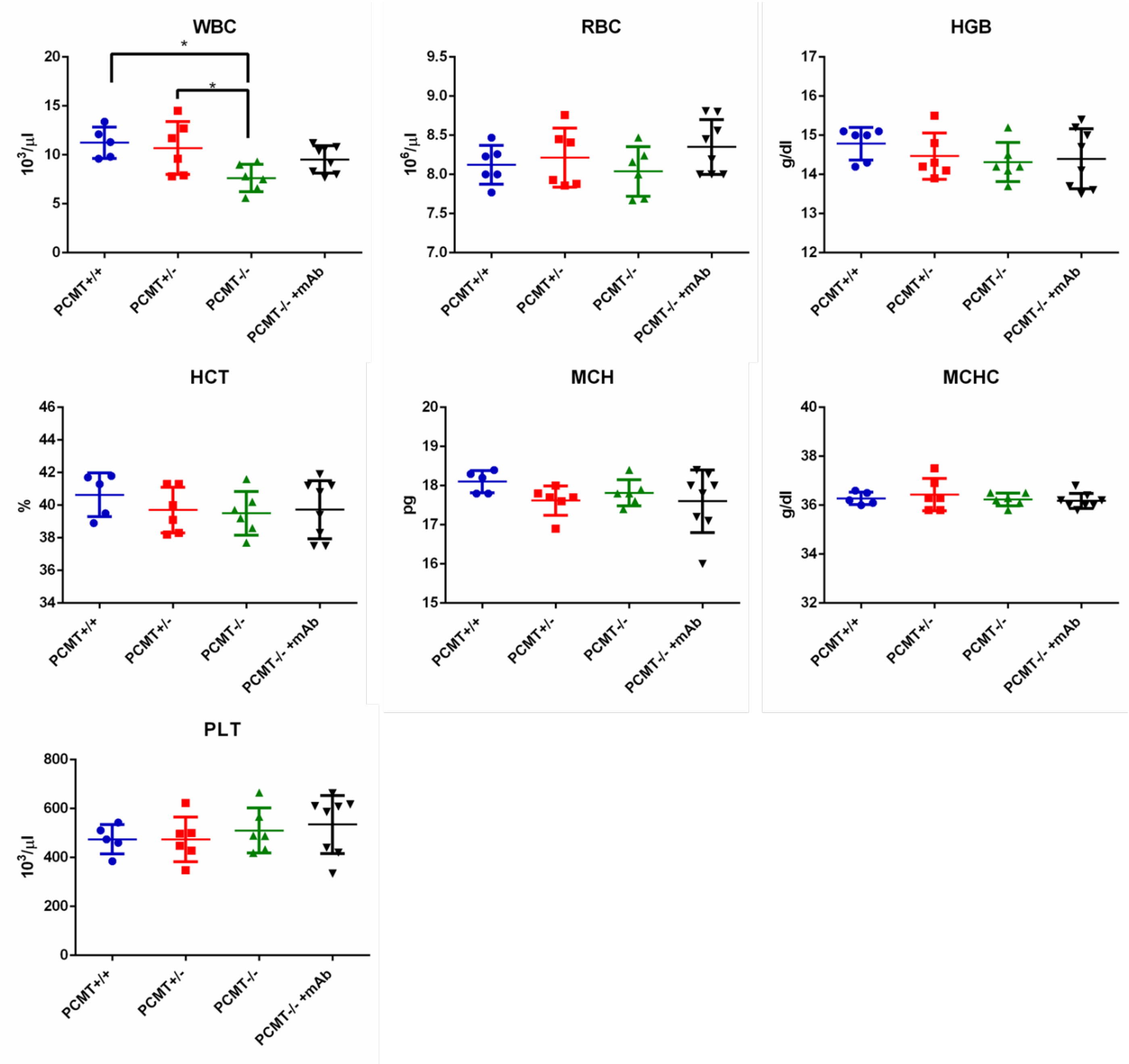


***Figure S3: Complete blood count is not altered in Pcmt1$^{-/-}$ mice***

*Total white blood cell (WBC) count was reduced in Pcmt1$^{-/-}$ mice, but red blood cells (RBC), haemoglobin level, oxygen transporters, and platelet numbers were not significantly altered relative to Pcmt1$^{+/+}$ mice. Data are shown as mean ± SEM (*p < 0.05) (n=6-8).*

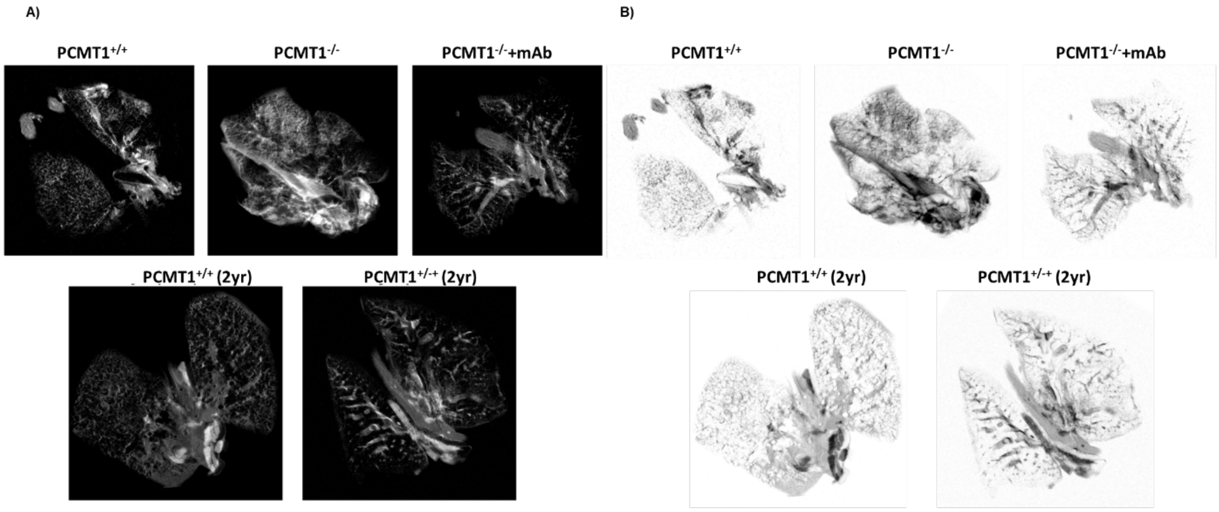


***Figure S4. MRI images of Pcmt1$^{+/+}$, Pcmt1$^{+/-}$, Pcmt1$^{-/-}$, and mAb-treated Pcmt1$^{-/-}$ lungs***
*Lung MRI representative contrast **(A)** and inverted **(B)** images for all genotypes assessed at age 5-6 weeks and 2 years (n=3).*

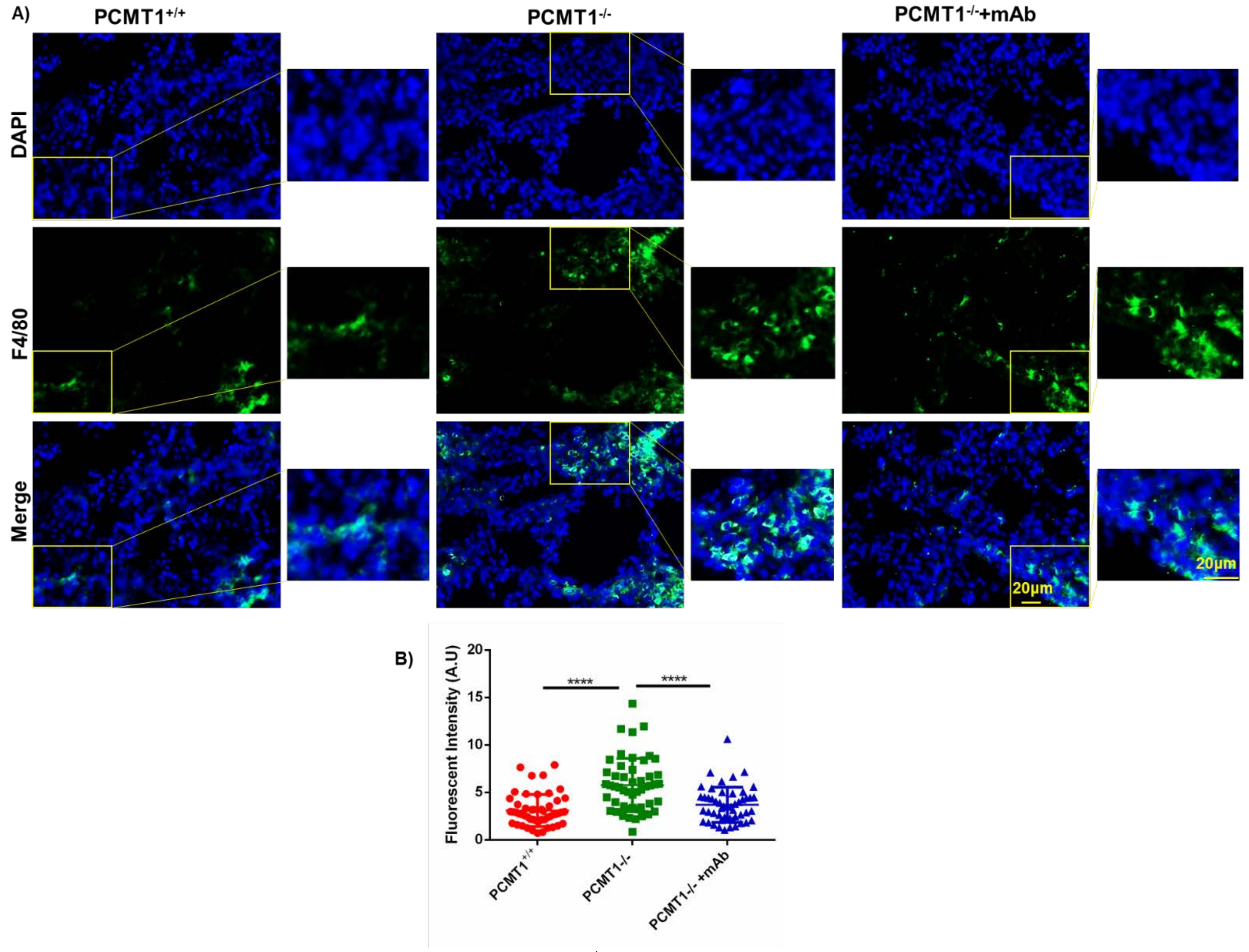


***Figure S5: Increased F4/80+ cell infiltration of Pcmt1$^{-/-}$ mouse lung***

*A) Representative immunostaining of F4/80+ macrophages in cryosectioned lung tissue from Pcmt1$^{+/+}$, Pcmt1$^{-/-}$, and mAb-treated Pcmt1$^{-/-}$ mice at age 5-6 weeks (n=5-7). (B) Graph represents fluorescence intensity of F4/80+ in cryosectioned lung tissue from Pcmt1$^{+/+}$, Pcmt1$^{-/-}$, and mAb-treated Pcmt1$^{-/-}$ mice. Results are shown as mean values ± SEM (**** $p<0.001$).*

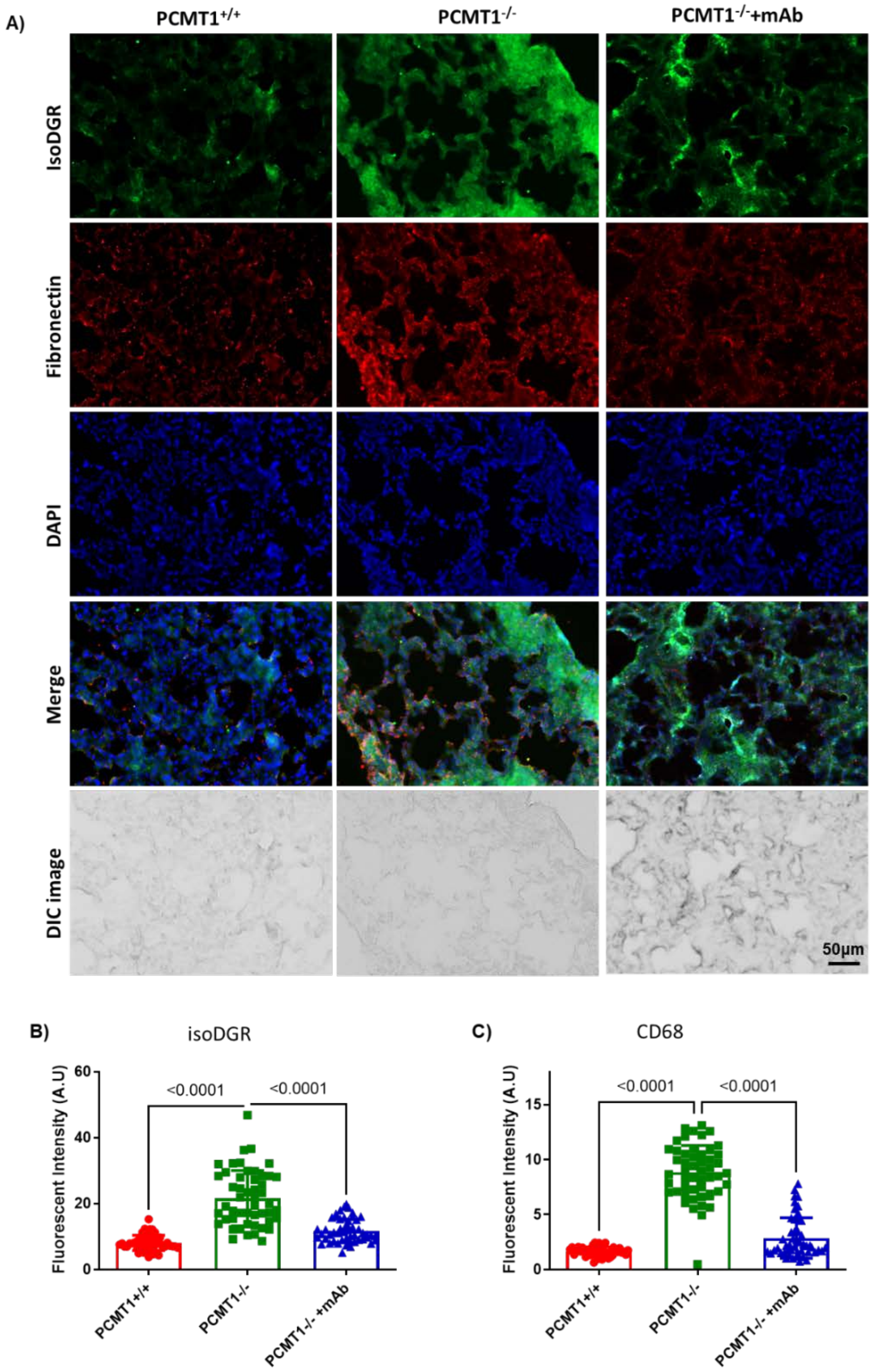


***Figure S6: IsoDGR accumulation and correlation with fibronectin in* Pcmt1$^{-/-}$ *lung tissue***

*A) Representative immunostaining of isoDGR protein distribution and correlation with fibronectin in cryosectioned lung tissue from Pcmt1$^{+/+}$, Pcmt1$^{-/-}$, and mAb-treated Pcmt1$^{-/-}$ mice at age 5-6 weeks (n=3). IsoDGR **(B)** and fibronectin **(C)** were quantified in Image J using 50 randomized regions in 5 images from 5 independent lung sections for each genotype (graphs display mean values for the same region from 5 images). Results are mean ± SEM (***p<0.001, **p<0.01, *p<0.05).*

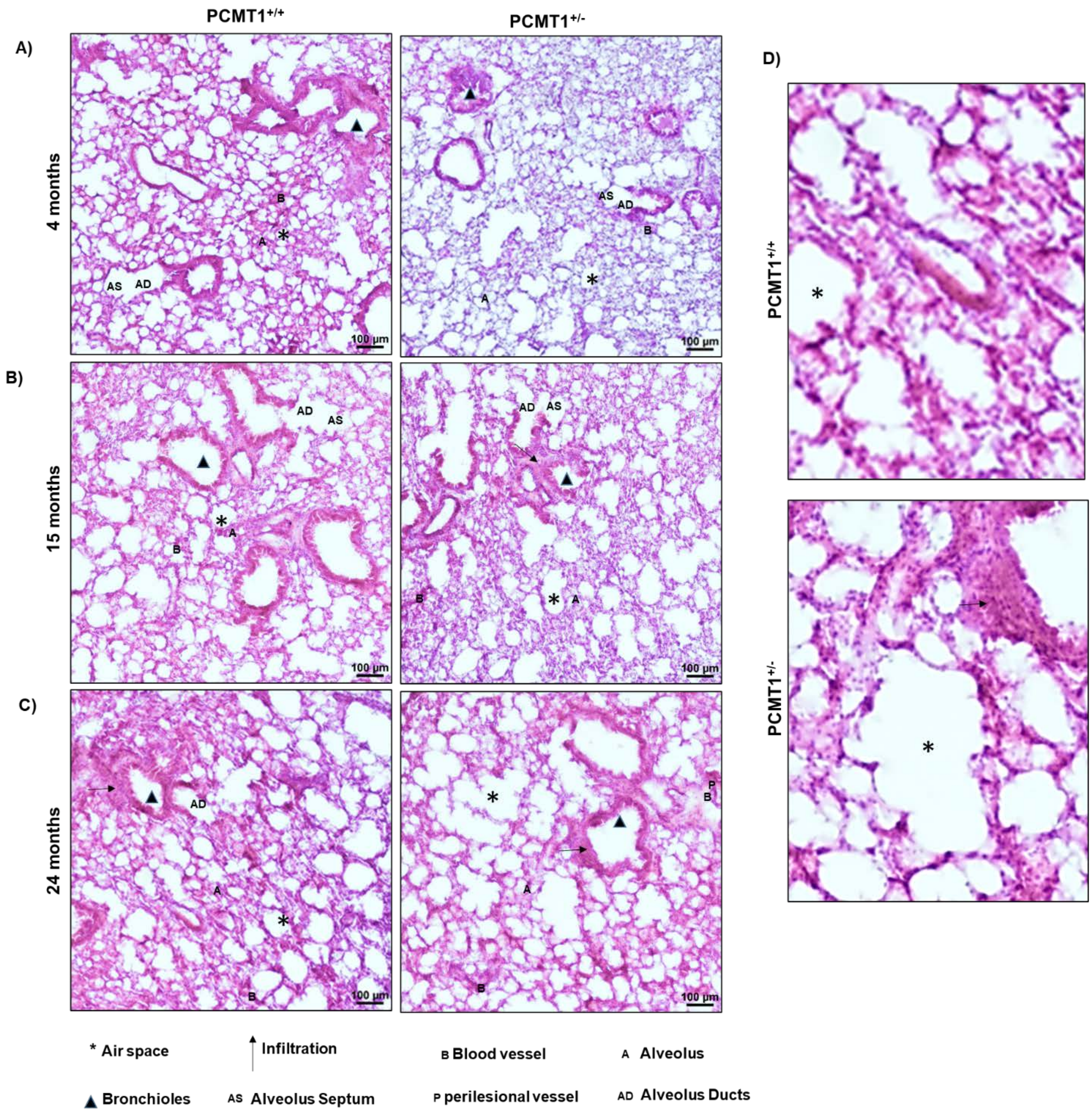


***Figure S7: Age-dependent airspace enlargement in lungs from Pcmt1$^{+/-}$ mice***

*Representative images of H&E-stained lung sections from Pcmt1$^{+/+}$ and Pcmt1$^{+/-}$ mice assessed at 4 months **(A)**, 15 months **(B),** or 24 months **(C)**. Magnified H&E images show lungs from Pcmt1$^{+/+}$ or Pcmt1$^{+/-}$ mice with air space enlargement and immune cell infiltration **(D)**.*

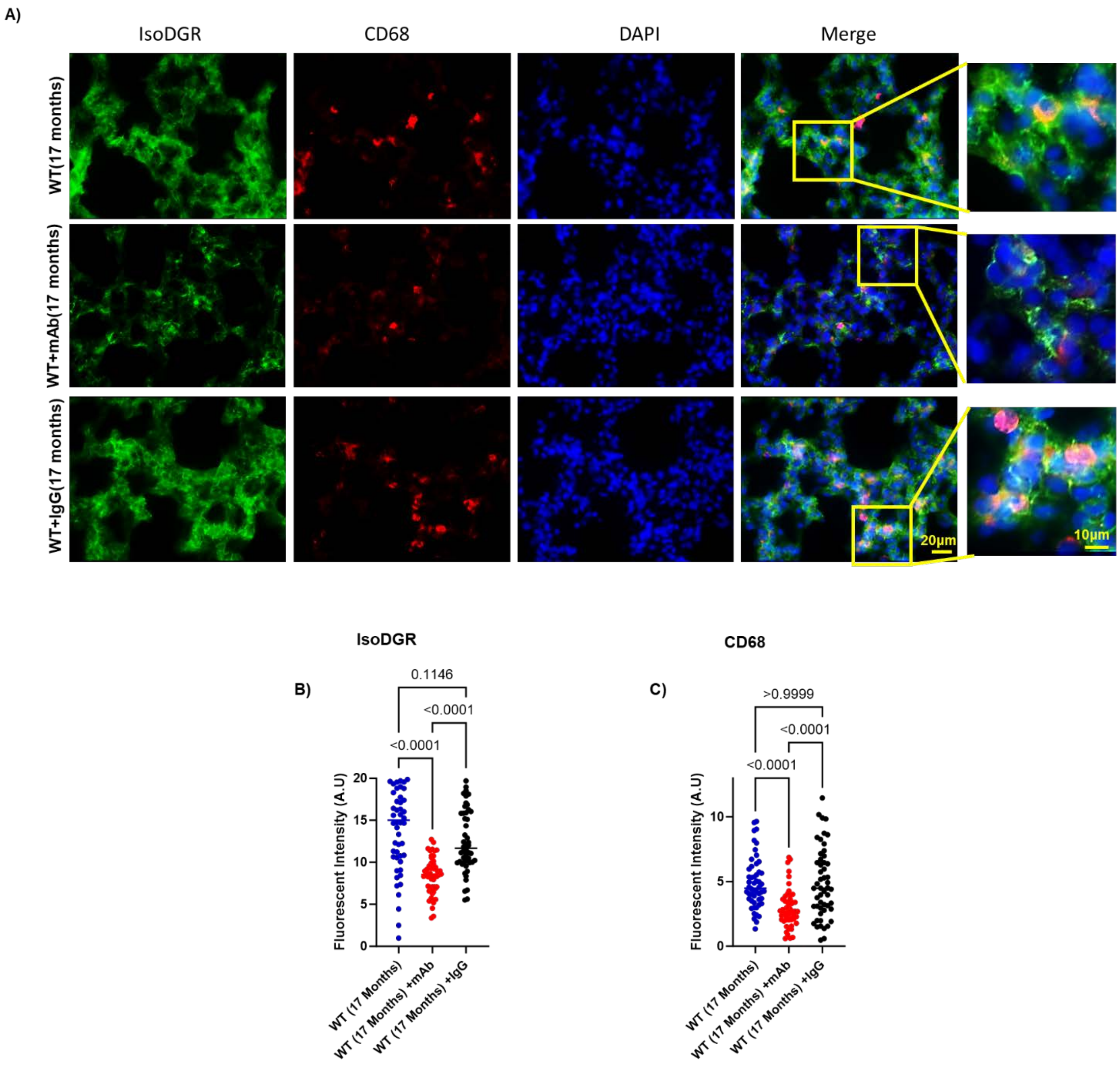


***Figure S8: Immune clearance of isoDGR-modified proteins in lung tissues of naturally-aged mice***

***(A)*** *Representative immunostaining images showing that isoDGR-proteins positive correlation with CD68*$^{+}$ *macrophages in cryosectioned lung tissues of naturally-aged WT mice from treated mice (compared with isotype IgG-injected controls)(n=5).*

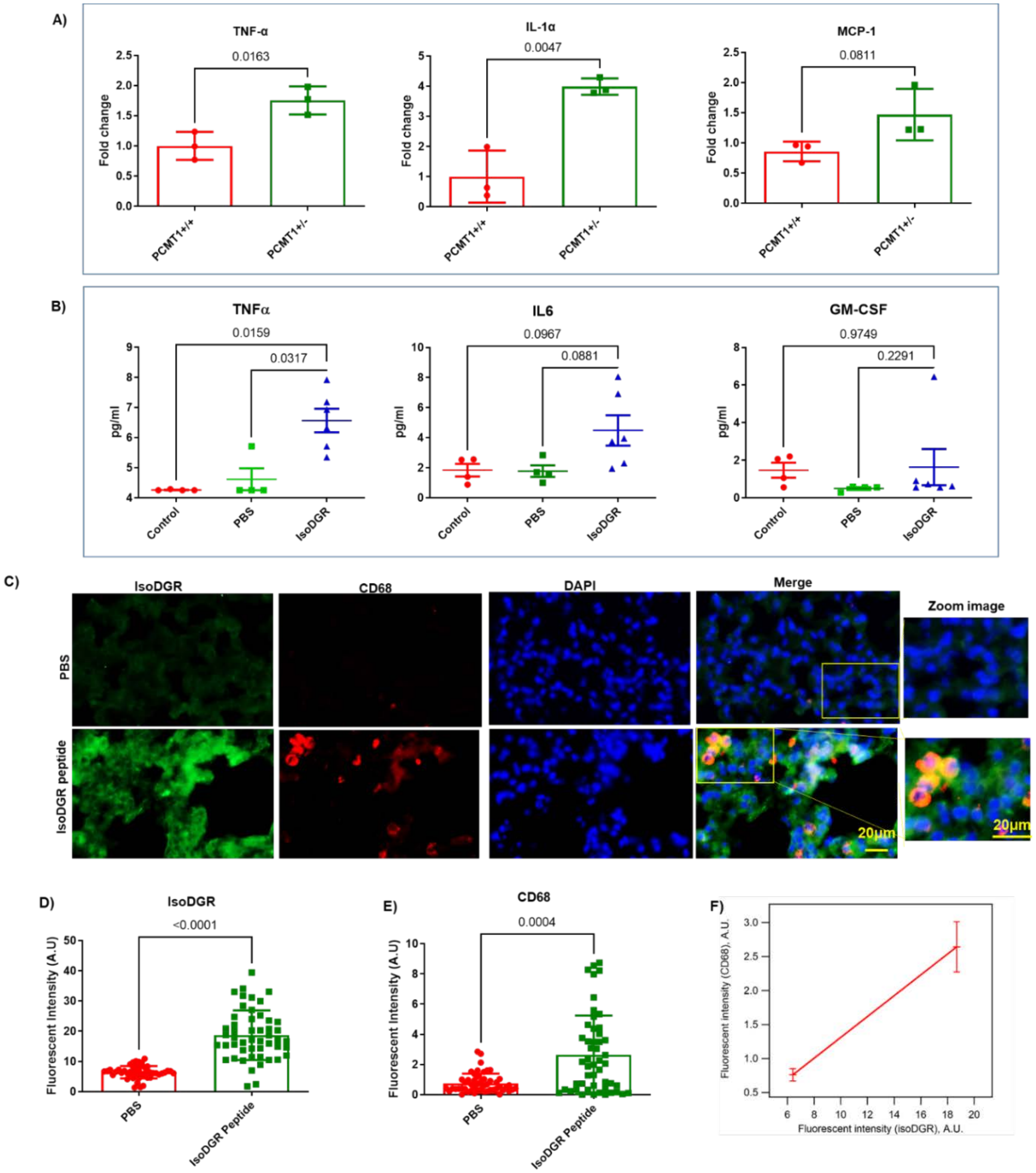


***Figure S9. Synthetic* isoDGR-peptides induce lung inflammation that resembles natural aging**

***(A)** Graph shows quantitation of pro- and anti-inflammatory cytokine levels in lungs from 2 year-old Pcmt1$^{+/+}$ and Pcmt1$^{+/-}$ mice (n=3). **(B)** Concentrations of MCP1, IL-6, and GM-CSF in lung interstitial fluids from Pcmt1$^{+/+}$ mice treated with synthetic isoDGR-peptide (n=6), or PBS-only vehicle control (n=4), or left untreated (n=4). **(C)** Representative immunostaining images showing that residual isoDGR peptide distribution co-localizes with CD68$^{+}$ macrophages in cryosectioned lung tissue from treated mice (compared with PBS-injected Pcmt1$^{+/+}$ controls)(n=4). Quantitative analysis of isoDGR-motif **(D)** and CD68 staining **(E)** in lung tissues. **(F)** CD68$^{+}$ macrophage infiltration of lung tissues was positively correlated with isoDGR levels. Results shown are mean ± SEM.*

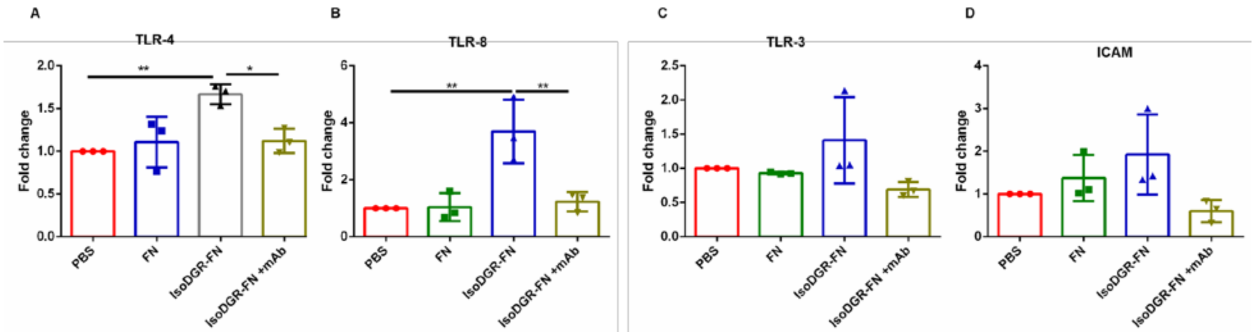


***Figure S10: IsoDGR-modified fibronectin activates TLR pathways** Graph showing quantitative PCR analysis of TLR-associated gene expression in HUVEC cells cultured with FN or isoDGR-FN in the presence or absence of motif-specific mAb (or PBS-only control); TLR4 **(A)**, TLR8 **(B)**, TLR3 **(C)**, ICAM **(D)**. Expression of GAPDH was used to normalize data (n=3). Results shown are mean values ± SEM (** p<0.01).*

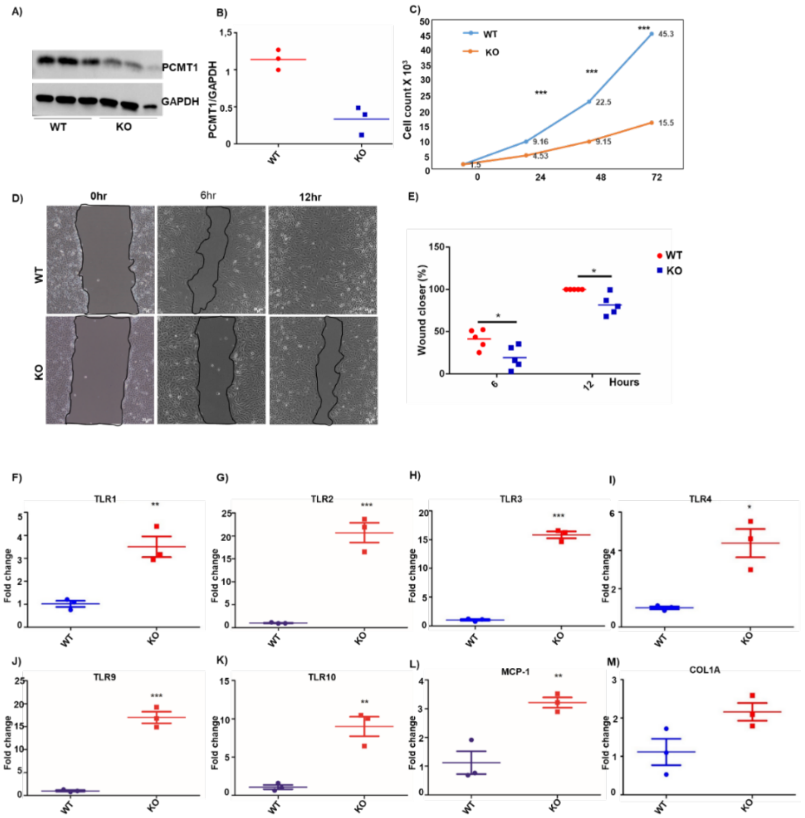


*Figure S11: **Pcmt1 knock-down impairs HUVEC proliferation / migration and activates TLR pathways. (A, B)** Protein lysates from HUVECs and HUVECsPcmt1 $^{KD}$ were subjected to western blot analysis using antibodies against Pcmt1 and GAPDH loading control (n=3). Results shown are mean values ± SD (***p≤0.001). **C**) Graph showing proliferation rates of Pcmt1$^{+/+}$ and Pcmt1-KO cells at 0, 24, 48 and 72h. (n=5) **D**) Representative images showing migration of HUVEC and HUVECPcmt1$^{KD}$ cells (n=5) **E**) Graph in right panel shows quantitation of migration rate comparing HUVEC and HUVECPcmt1$^{KD}$ (mean ± SD, n=3). **F-M**) Graphs show quantitative PCR analysis of TLR pathway-associated gene expression in HUVECs and HUVECsPcmt1 $^{KD}$. Data are normalised to GAPDH expression (n=3). Results are shown as mean values ± SEM (***p<0.001, **p<0.01, *p<0.05).*

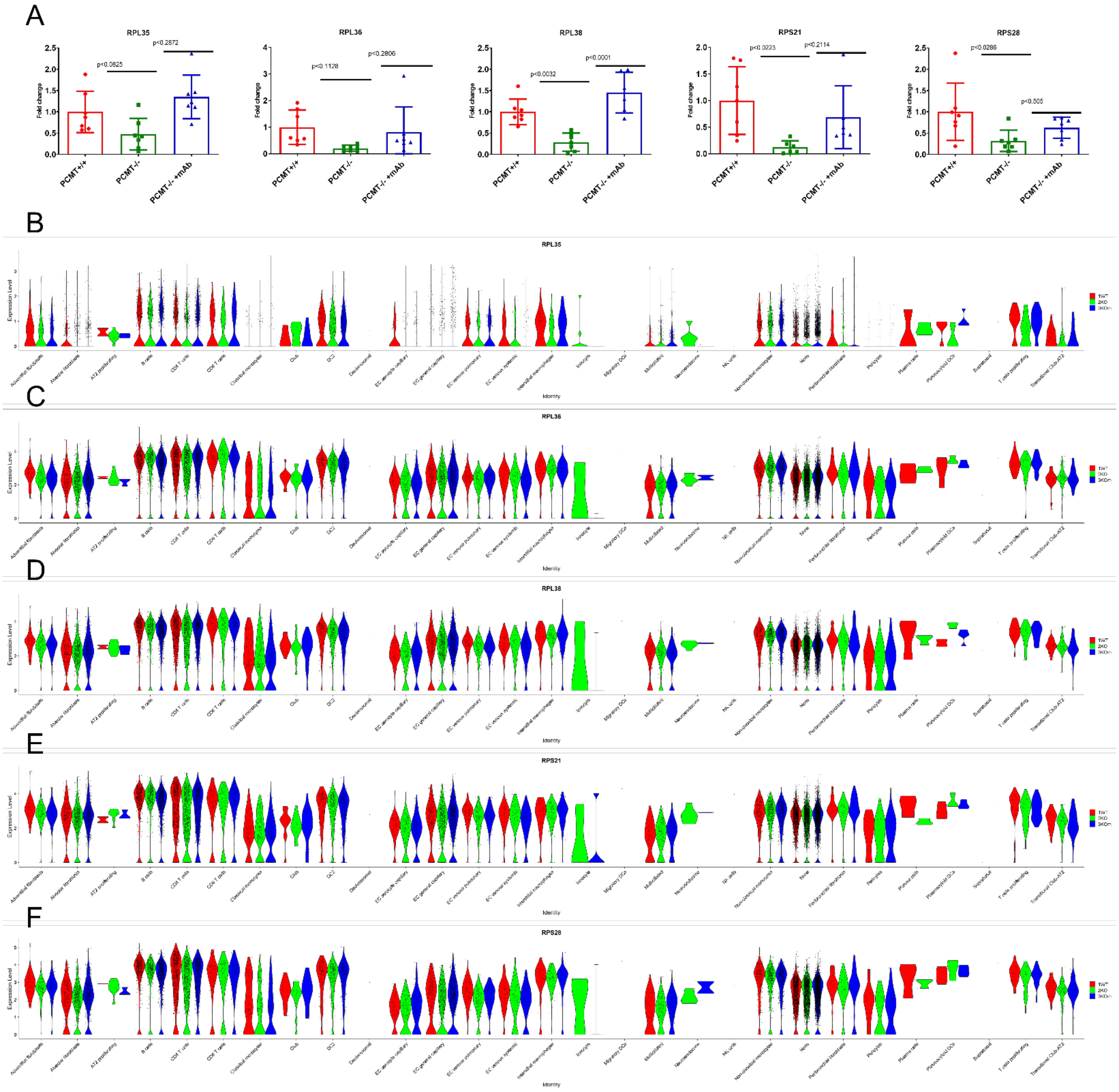


***Figure S12: IsoDGR mAb treatment increases expression of ribosomal synthetic machinery proteins in Pcmt$^{-/-}$ mouse lung.*** *(A) Graph showing quantitative PCR analysis of ribosome-associated genes RPL35, RPL36, RPL38, RPS21, and RPS28 in lung tissues from Pcmt1$^{+/+}$, Pcmt1$^{-/-}$ and mAb-treated Pcmt1$^{-/}$ mice at 5-6 weeks. Expression of GAPDH was used to normalize data. Results are mean ± SEM (n=6). Violin plots of (B) RPL35, (C) RPL36, (D) RPL38, (E) RPS21, and (F) RPS28 expression across various lung cell types, as determined by scRNA-seq analysis.*

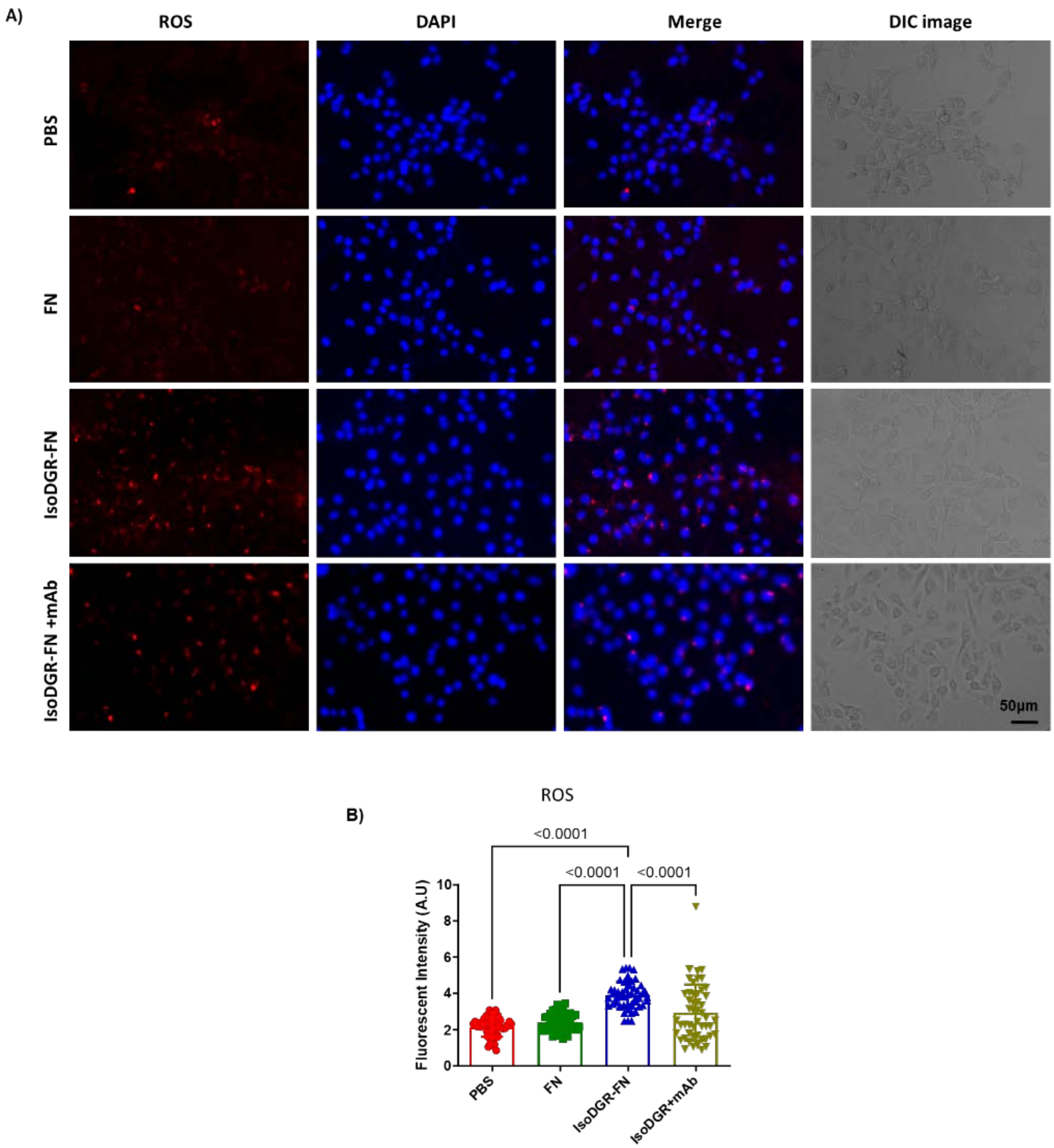


***Figure S13: IsoDGR-modified fibronectin increases ROS production in*** **HULEC-5a cells**

*(A) Representative immunostaining images showing ROS distribution in HULEC-5a cells cultured with PBS only, native FN, or isoDGR-FN, either in the presence or absence of anti-isoDGR mAb.* ***(B)*** *ROS fluorescence was quantified in Image J using 50 randomized regions from 5 images of 5 independent experiments (graphs show average values for the same region from 5 images). Results shown are mean values ± SEM.*

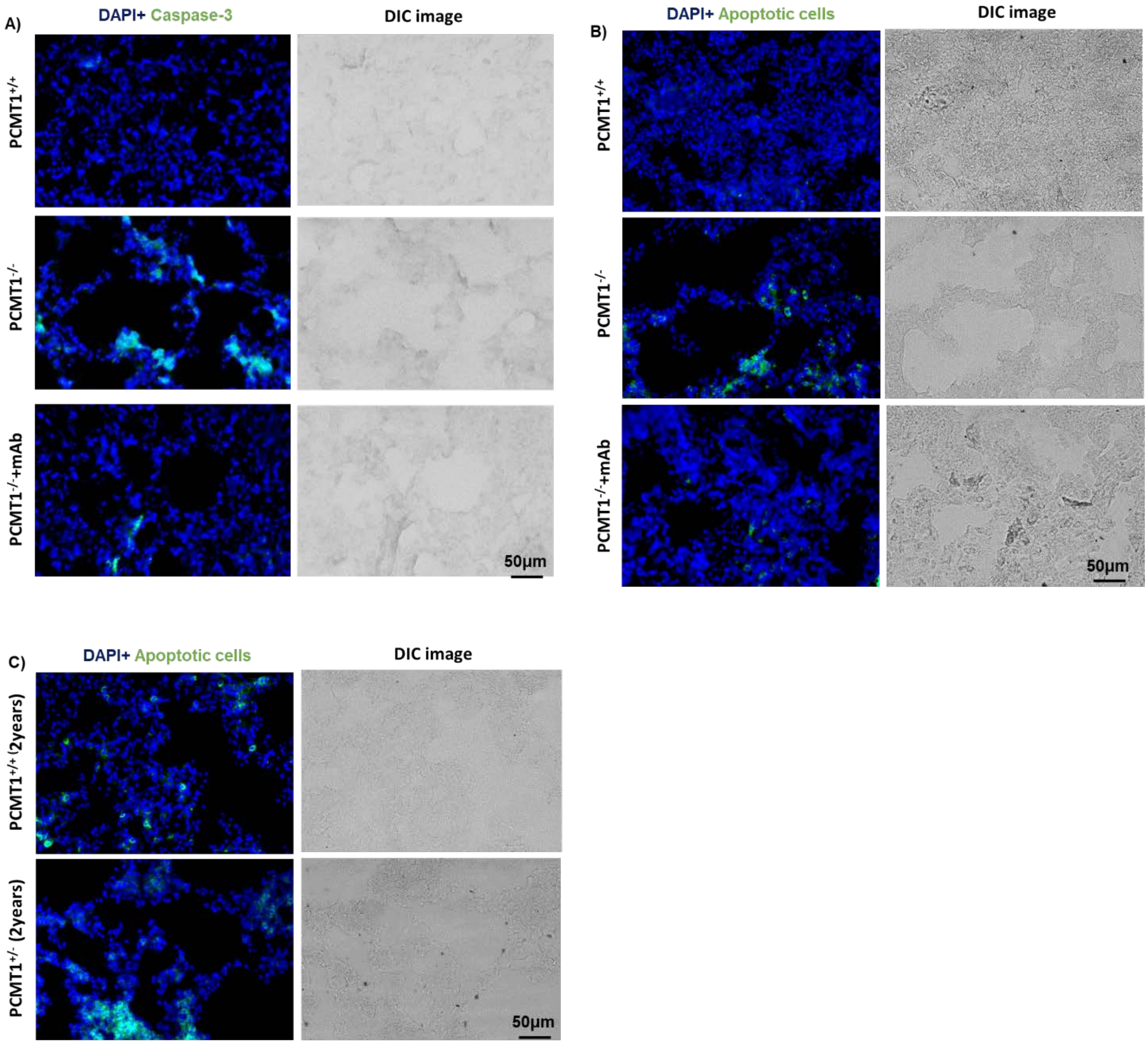


***Figure S14: IsoDGR-modified proteins induce apoptosis of lung parenchymal and EC cells***

*Representative images of Tunel assay identifying caspase-3 positive cells **(A)** and apoptotic cells (**B**) in lung tissue from 5-6 weeks old Pcmt1$^{+/+}$, Pcmt1$^{-/-}$ and mAb-treated Pcmt1$^{-/-}$ mice, or **(C)** in lung sections from 2-year-old Pcmt1$^{+/+}$ and Pcmt1$^{+/-}$ mice (n=3 per group).*

**Table S1 Primers used in genotyping mouse Pcmt1 gene.**

| Primer Name | Genotyping Primer |
|---|---|
| oIMR108 | CGGCTGCATACGCTTGATC |
| oIMR1081 | CGACAAGACCGGCTTCCAT |
| oIMR1544 | CACGTGGGCTCCAGCATT |
| oIMR3580 | TCACCAGTCATTTCTGCCTTTG |

**Table S2 RTPCR Primers for mouse tissue**

| Gene name | Primer sequences |
|---|---|
| **RPS28-F** | GCCGCTCTATCATCCGAAAT |
| **RPS28-R** | GCTGCAAGATTAACGCAACC |
| **RPL38-F** | GGTACCTTTACACCCTGGTTATC |
| **RPL38-R** | CAGAGGGCTGGTTCATTTCA |
| **RPL36-F** | AAACGTCAGTAAGCCGAGAC |
| **RPL36-R** | TGGACACTTTGAGCAACTCC |
| **RPL35A-F** | GCTGGGAACAGGACTTCTAAC |
| **RPL35A-R** | CCGTTTCATCTCGGGCATAA |
| **RPS21-F** | ACCACAGGCCGGTTTAATG |
| **RPS21-R** | GCCTTAGCCAATCGGAGAATAG |
| **TLR1-F** | GCTGGTGTTAGGAGATGCTTAT |
| **TLR1-R** | GACGGACACATCCAGAAGAAA |
| **TLR3-F** | GTGCATCGGATTCTTGGTTTC |
| **TLR3-R** | GACCCAGTCTCTGTCTTTATGG |
| **TLR4-F** | AGTATCGAGAGGCTCAGGTATAG |
| **TLR4-R** | TACAGGATGCAGGACAAGTAATC |
| **TLR8-F** | CCTCTCTAAGGCTAGGGTAACT |
| **TLR8-R** | TGCCCAGAAGACAGCATTT |
| **TIRAP-F** | CCATCATAGAGGTGGCTTTCC |

| | |
|---|---|
| **TIRAP-R** | TCGATGCCTTTATCTGCTACTG |
| **TRAF6-F** | GGTACCTGAGAGATGCCTAAAC |
| **TRAF6-R** | GACAGTCACCTCTCCACATTAG |
| **FADD-F** | CCTTTGGCTGGGTGTGATAATA |
| **FADD-R** | TCTCTATGTAGGCTAGGCTGTC |
| **MCP1-F** | TGATCCCAATGAGTAGGCTGGAG |
| **MCP1-R** | ATGTCTGGACCCATTCCTTCTTG |
| **IL-1a-F** | CAAACTGATGAAGCTCGTCA |
| **IL-1a-R** | TCTCCTTGAGCGCTCACGAA |
| **IL-8-F** | CGTGGCTCTCTTGGCAGCCTTC |
| **IL-8-R** | TCCACAACCCTCTGCACCCAGTT |
| **GAPDH-R** | AAGATGGTGATGGGCTTCCCG |
| **GAPDH-F** | TGGCAAAGTGGAGATTGTTGCC |
| **IL-3-F** | TGAAGGACCCTCTCTGAGGA |
| **IL-3-R** | CGCAGATCATTCGCAGAT |
| **CCL4-F** | CAAACCTAACCCCGAGCAACAC |
| **CCL4-R** | GGTCTCATAGTAATCCATCACAAAGC |
| **IL-10-F** | ATGCAGGACTTTAAGGGTTACTTGGGT |
| **IL-10-R** | ATTTCGGAGAGAGGTACAAACGAGGTTT |
| **IL-12p40-F** | CAGAAGCTAACCATCTCCTGGTTTG |
| **IL-12p40-R** | TCCGGAGTAATTTGGTGCTTCACAC |
| **TNF-α-F** | GCCTCTTCTCATTCCTGCTTG |
| **TNF-α-R** | CTGATGAGAGGGAGGCCATT |
| **TLR9-F** | TGGTTACCTGGCAAGACGC |
| **TLR9-R** | GGAAACTGGCACGCAAGAG |
| **IRAK-4-F** | CCATCGTGGCGGTGAAG |
| **IRAK-4-R** | GTGCTGACACGTTGCCATTACT |

| | |
|---|---|
| **mt-ND1-F** | GCTTTACGAGCCGTAGCCCA |
| **mtND1-R** | GGGTCAGGCTGGCAGAAGTAA |
| **mt-ND2-F** | CCTCCTGGCCATCGTACTCA |
| **mt-ND2-R** | GAATGGGGCGAGGCCTAGTT |
| **mt-ND3-F** | TAGTTGCATTCTGACTCCCCCA |
| **mt-ND3-R** | GAGAATGGTAGACGTGCAGAGC |
| **mt-ND4-F** | CGCCTACTCCTCAGTTAGCCA |
| **mt-ND4-R** | TGATGTGAGGCCATGTGCGA |
| **mt-ND4l-F** | AGCTCCATACCAATCCCCATCAC |
| **mt-ND4l-F** | AGCTCCATACCAATCCCCATCAC |
| **mt-ND5-F** | GGCCCTACACCAGTTTCAGC |
| **mt-ND5-R** | AGGGCTCCGAGGCAAAGTAT |
| **mt-ND6-F** | CTTGATGGTTTGGGAGATTGG |
| **mt-ND6-R** | ACCCGCAAACAAAGATCACC |
| **mt-Cytb-F** | TCCTTCATGTCGGACGAGGC |
| **mt-Cytb-R** | AATGCTGTGGCTATGACTGCG |
| **mt-CO1-F** | TCAACATGAAACCCCCAGCCA |
| **mt-CO1-R** | GCGGCTAGCACTGGTAGTGA |
| **mt-ATP6-F** | AGCTCACTTGCCCACTTCCT |
| **mt-ATP6-R** | AAGCCGGACTGCTAATGCCA |
| **mt-CO2-R** | TCCTAGGGAGGGGACTGCTC |
| **mt-CO2-F** | ACCTGGTGAACTACGACTGCT |

**Table S3 RT-PCR Primers for human cells**

| Gene Name | Primer sequence |
|---|---|
| **TLR1-F** | CATGGCCAGGAGGACTTATTT |
| **TLR1-R** | TGCTTGCTCTGTCAGCTTAATA |
| **TLR2-F** | GAAGAGTGAGTGGTGCAAGTAT |
| **TLR2-R** | AATGGGCTCCAGAAGAATGAG |
| **TLR3-F** | CCCTGGTGGTCCCATTTATTT |
| **TLR3-R** | CTCAACTGGGATCTCGTCAAAG |
| **TLR4-F** | GATGAGGACTGGGTAAGGAATG |
| **TLR4-R** | GGCCACACCGGGAATAAA |
| **TLR8-F** | CACCAGAGACATAGGCATCAC |
| **TLR8-R** | TCGCATGGCTTACATGAGTATAG |
| **TLR9-F** | GCTAGACCTGTCCCACAATAAG |
| **TLR9-R** | AAAGGGCTGGCTGTTGTAG |
| **TLR10-F** | GCTAGACCTGTCCCACAATAAG |
| **TLR10-R** | AAAGGGCTGGCTGTTGTAG |
| **GAPDH-F** | GTGGTCTCCTCTGACTTCAACA |
| **GAPDH-R** | CTCTTCCTCTTGTGCTCTTGCT |
| **ICAM1-F** | CTCCAATGTGCCAGGCTTG |
| **ICAM1-R** | CAGTGGGAAAGTGCCATCCT |
| **MCP-1-F** | CAGATGCAATCAATGCCCCAG |
| **MCP-1-R** | ATAAAACAGGGTGTCTGGGGAAAGC |
| **COL1A1-F** | TCTGCGACAACGGCAAGGTG |
| **COL1A1-R** | GACGCCGGTGGTTTCTTGGT |
